# Brain mechanisms of mathematical development: A longitudinal fMRI study from preschool to second grade

**DOI:** 10.64898/2026.01.19.700370

**Authors:** Théo Morfoisse, Severine Becuwe, Marie Palu, Cassandra Potier-Watkins, Ghislaine Dehaene-Lambertz, Stanislas Dehaene

## Abstract

How does the developing brain, initially equipped only with elementary mathematical intuitions, acquire higher mathematical concepts? Through a longitudinal functional MRI study of children from preschool through first and second grade, we tracked how neural responses to mathematical and non-mathematical statements change in the first two years of formal schooling and use the data to evaluate several theories of developmental change. Before school, when listening to math statements, children already engage an adult-like cortical network, with partial specialization for geometry. Over the first two years of school, we observe an overall increase in math-related activation, a small recruitment of additional neural territory, reduced activation for facts that get better known, and a small overall increase in the dimensionality of representational space. fMRI responses to individual sentences suggest that these mechanisms, particularly in left inferior frontal gyrus and bilateral intraparietal sulcus, all contribute to children’s growing mastery of mathematical concepts.

## Introduction

While the foundations of mathematics are intuitive and accessible to all, its scope quickly widens to include abstract, complex and education-dependent concepts. Proto-mathematical intuitions – such as approximate number and non-symbolic geometry - emerge early in life (*1–4*), are shared across cultures (*5–8*), and with other species such as monkeys (*9*, *10*), fishes (*11*, *12*) or even bees (*13*, *14*), suggesting a phylogenetically ancient foundation for non-symbolic mathematical cognition (*15*, *16*). In contrast, formal symbolic mathematics is unique to humans (*8*, *17*), rests on years of cultural transmission and structured education (*18*, *19*) and remains a source of difficulty for many children (*20*). How do children overcome the limits of their intuitive non-verbal mathematical abilities and expand their knowledge to include symbolic mathematics, and how does the brain adapt to this learning? While the neural mechanisms of reading acquisition are increasingly understood (*21*), how the brain changes with math education remains largely unknown.

One prevailing theory suggests that symbolic mathematics repurpose evolutionary ancient brain circuits originally underlying proto-mathematical skills (*15*, *17*, *22*, *23*). Indeed, neuroimaging studies have identified a reproducible math-responsive cortical network involving bilateral dorsal prefrontal, intraparietal, and ventrolateral inferior temporal areas in adults (*15*, *24*, *25*) which closely resembles the network already observed in infants (*1*, *26*), children (*27*, *28*) and non-human primates (*16*, *29*). This network supports not only non-symbolic or elementary mathematics (*30–35*), but also symbolic arithmetic (*32*, *33*, *35–41*), and higher-order mathematical reasoning (*42–46*). Even professional mathematicians continue to engage the same overall areas when processing advanced mathematical concepts (*22*, *42*). Yet, despite this anatomical continuity, it remains unclear how the same neural circuits can evolve from supporting basic numerical and geometric intuitions to enabling the school-based acquisition of complex symbolic mathematics. Two complementary lines of research have begun to shed light on this issue. A first series of studies comparing children to adults (*30–33*, *36*, *39*, *40*, *43*, *47*, *48*) notably revealed a developmental shift from a strong reliance on frontal regions in children to greater involvement of parietal areas in adults (*27*, *32*), likely reflecting an automatization of arithmetic fact retrieval. However, by contrasting only two points of the developmental continuum, such cross-sectional studies offer limited insight into the neural changes in between. A second set of studies has focused on finer-grained developmental trajectories, either through cross-sectional comparison of children at different ages (*35*, *37*, *38*, *41*, *43*, *49–51*), or through longitudinal designs (*39*, *52*, *53*). The latter, in particular, yielded important observations such as a transient increase in hippocampal activity and hippocampal-prefrontal-parietal functional correlation as children improve their memory for arithmetic facts (*53*), accompanied by an increased coupling between intraparietal and inferior temporal cortices and a decoupling from prefrontal cortex (*52*, *53*).

While these studies have begun to uncover neural correlates of mathematical development, significant limitations remain. First, most studies have focused on basic numerical tasks, with only a few exploring more complex or educationally relevant materials (*43*, *45*, *49*, *50*). Second, longitudinal brain-imaging studies remain scarce and often limited to small, heterogeneous samples (e.g., 7-14 year-olds in (*52*)). Third, the pivotal transition from preschool intuition to formal school-based instruction, is still largely unexplored (*31*, *35*, *39*, *43*). For all these reasons, we still lack a clear understanding of how the brain changes when children enter school and begin learning symbolic mathematics.

To address these gaps, we conducted a longitudinal study in children, with three consecutive functional MRI scans acquired one year apart (n = 149 scans) from the end of preschool through first and second grade – a pivotal period of rapid cognitive and educational change. Children performed a sentence listening task (*22*, *42*) evaluating the veracity of statements in three domains: mathematics (including arithmetic and geometry), general knowledge and social concepts. Unlike previous studies focusing on non-symbolic numerosity or basic numerals understanding, this design targeted a broader range of mathematical vocabulary and, crucially, their composition into meaningful but possibly incorrect statements.

Our design probed mathematical knowledge at three complementary levels: (1) domain-level differences, by comparing mathematical with non-mathematical statements; (2) subdomain specialization within mathematics, by contrasting arithmetic and geometry; and (3) fine-grained representational structure, by mapping neural responses to individual sentences thus characterizing the emerging space of mathematical concepts. Specifically, we first characterized the children’s mathematical network by contrasting mathematical statements with those from the two control domains. We then investigated developmental changes using domain-specific regions of interest (ROIs) defined from adult fMRI data acquired with a similar paradigm (*54*), allowing us to test several prominent hypotheses regarding brain changes underlying child development (*55*) (figure 1):

- **Early cortical organization** (figure 1A). Upon school entry, does the child’s brain already route verbal statements depending on their contents into adult-like circuits more responsive to math than to non-math semantic knowledge? This hypothesis is rooted in the cortical recycling model and supported by limited prior data in preschoolers and infants (*26*, *31*, *43*). Alternatively, some of the areas might still be immature, poorly segregated, or unresponsive to composite math statements. Empiricist neo-connectionist theories (e.g. *56*) and parallels between human development and learning in large-language models (e.g. *57*) should predict that a long exposure to verbal inputs is required before a segregation of math and non-math sentences emerges. As a finer test, we contrasted arithmetic and geometry, which also exhibit a partial segregation in adults (*42*, *58*), to examine whether, at this age, the math-responsive network is still functionally unified, or already fully differentiated into specialized sub-networks.
- **Developmental change in cortical space** (figure 1B). How does cortical activity in the math network evolve during development? Does it involve an activation increase, as expected if the recycled areas become increasingly responsive to words and symbols (*42*); a decrease, if neural representations become sparser, more sharply tuned (*59*) and/or mobilize fewer cognitive resources (*55*); or both, i.e. a prefrontal decrease and a parietal increase due to greater reliance on automatized posterior circuits, as suggested by several previous experiments (*27*, *32*, *60*, *61*)? If an increase is observed, does it merely involve a strengthening of existing neural responses, as observed when monkeys were trained for numerosity (*62*) or an expansion into new cortical territory, as observed in the case of reading (*21*, *63*)? And can the effects of age, school exposure, and individual proficiency on these changes be separated?
- **Developmental change in high-dimensional neural space** (figure 1C). Assuming that words, and sentences are represented as neural vectors embedded in a high-dimensional manifold (*64–67*), how does this manifold change as new concepts are understood? Beyond conventional subject-level analyses, we adopted a sentence-centered approach that treats each sentence as a unit of analysis, allowing us to quantify the dimensionality of their neural representational space. We hypothesized that dimensionality would increase with mathematical development, reflecting the progressive differentiation of a growing number of conceptual expressions within a shared neural network (*68*). Alternatively, dimensionality might remain stable, or even decrease, if newly acquired concepts become aligned with pre-existing neural representations.

**Figure 1.**
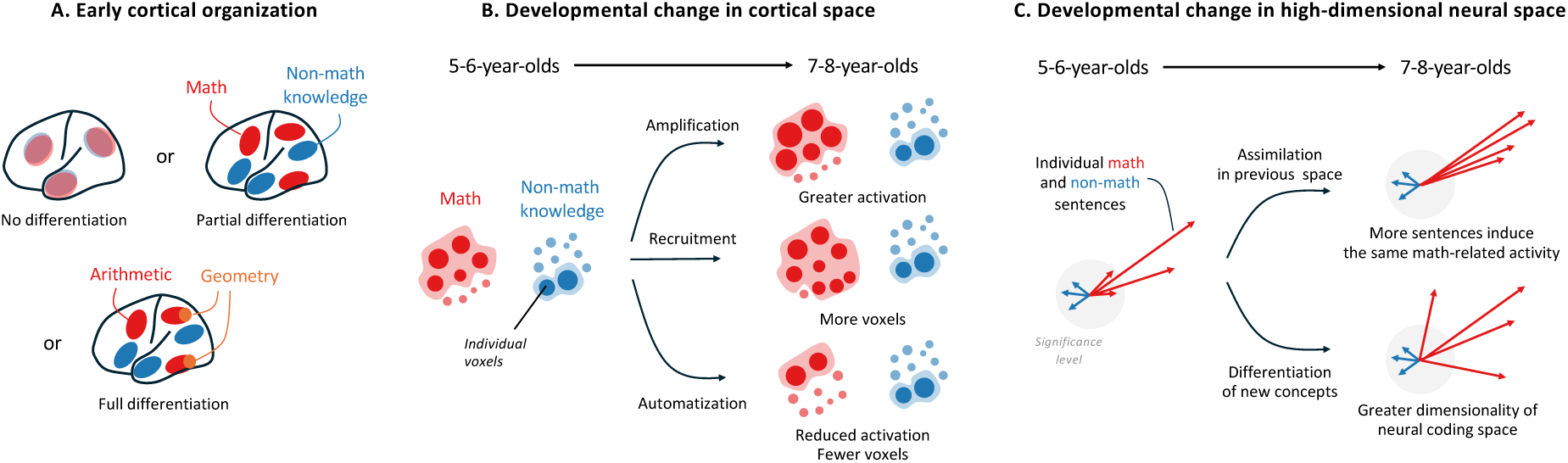
Open issues concerning the neural correlates of mathematical education. (A) Early cortical organization. Prior to schooling, semantic domains of math and non-math may show minimal, partial, or full separation at the cortical level. **(B) Developmental change in cortical space.** Each circle schematizes a voxel, whose diameter indicates its activation in the contrast of math vs non-math statements (respectively red vs blue). Across the first two years of schooling, voxels may follow (1) an *amplification* model, where selective voxels become increasingly activated over time; (2) a *recruitment* model, where previously inactive or non-selective voxels are recruited for math; and (3) an *automatization* model, where activation to the same statements diminishes as development progresses. **(C) Developmental change in high-dimensional neural space.** Each vector represents the neural activation to each math or non-math statement in high-dimensional neural space. With development, sentence-level activation may follow (1) an *assimilation* model, where newly understood statements induce the same vector of math-related activity as previous ones; or (2) a *differentiation* model, where statements get represented by increasingly distinct neural vectors, resulting in an increased dimensionality of neural coding space.

## Results

### Behavior

Children performed significantly above chance across all conditions, regardless of the testing period (p ≤ .001, figure S1A). Binomial mixed-model regression, with condition and age as fixed effects, and participant as a random effect revealed that children were significantly more accurate in judging the truth of general knowledge (z = - 4.08, p < .001) and social sentences (z = - 5.29, p < .001) compared to mathematical ones, and that errors decreased with age across all three conditions (z = - 3.91, *p < .001*), with no significant interactions between age and condition (table S1). Similar mixed-model regressions on reaction times, revealed that children responded more quickly to social (t = −6.29, p < .001) and general knowledge (t = −6.63, p < .001) sentences than to mathematical ones (table S2). No significant effect of age on reaction times was observed (t = 0.87, p = .38), but a significant interaction indicated that reaction times for social sentences decreased more with age than those for mathematical sentences (t = −2.13, p = .033) (figure S1B).

### fMRI characterization of the math-responsive network in 5-to 8-year-olds

Actively listening to mathematical statements, as opposed to non-math sentences (GK and Social) activated a network of bilateral brain regions in 5-to 8-year-old children, including the intraparietal sulci (*IPS*), inferior temporal gyri (*ITG*), inferior frontal gyri (*IFG*), middle frontal gyri (*MFG*) and posterior cingulate gyri (figure 2A). Importantly, this math-responsive network was identified at all three ages tested (figure 2A, bottom row). Restricting the analysis to correct trials or controlling for trial-by-trial reaction time yielded virtually identical results (figure S3). As in adults, there was a double dissociation with other brain areas showing a greater response to social sentences, including the bilateral temporal and angular gyri, left inferior frontal gyrus, medial frontal gyrus and precuneus (figure S5). For direct comparison, figure S4A-B shows the math-responsive and social-responsive brain regions identified in an adult cohort using a similar paradigm(*54*).

**Figure 2.**
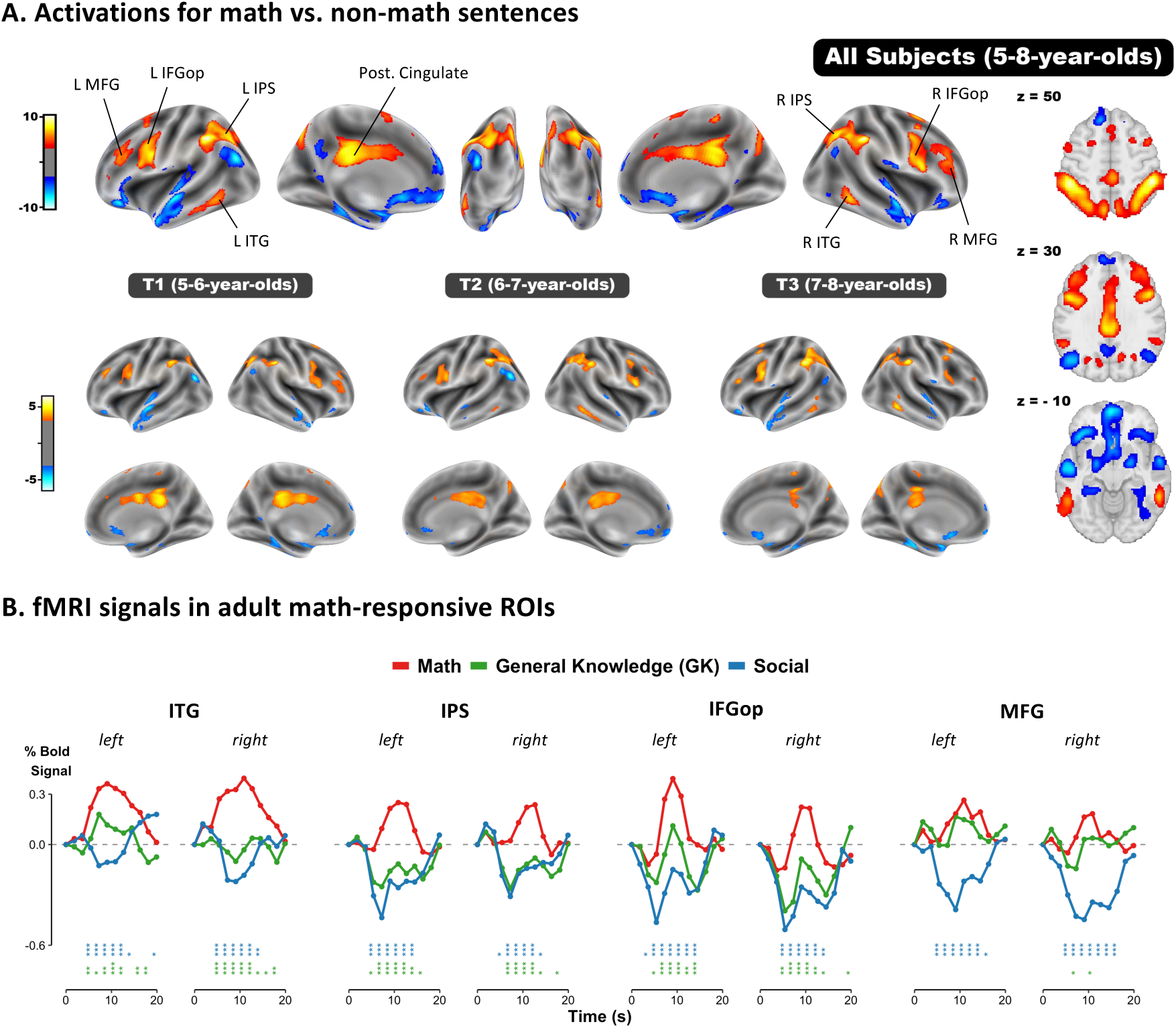
An adult-like math-responsive network in young children. **(A)** fMRI contrast between math and non-math sentences. The top row shows the joint analysis of all data, while bottom plots show separately the first fMRI (T1, 5-6-year-olds), second fMRI (T2, 6-7-year-olds), and third fMRI (T3, 7-8-year-olds). Voxel-wise p < 0.001, FDR-corrected α < 0.05. **(B)** Time course of average fMRI signals in math-related ROIs extracted from an adult cohort using a similar paradigm(*54*). Stars indicate the significance of independent mixed-model regressions conducted at each time step (TR), with condition as fixed effect and participant as random effect (*•p < 0.1; *p < 0.05; **p < 0.01; ***p < 0.001*). Blue stars: Math vs. Social. Green stars: Math vs. GK (general knowledge).

From these adult activation maps, we defined eight math-responsive ROIs, specifically the left and right ITG, IPS, IFGop and MFG (see Methods) and computed the fMRI time course signal for all three conditions within each of these regions for each child (figure 2B). Linear mixed-model regressions performed at each time step (TR), revealed that math sentences elicited significantly stronger responses compared to social sentences in all regions, with differences emerging around 5-7 seconds after sentence onset (p < .001 and t > 4 in all regions for TR = 4). A similar divergence was observed for math vs. general knowledge sentences, in all but two regions - the left and right MFG. Importantly, similar results were observed when we restricted the analysis to preschoolers or children within one month of entering first grade (figure S6).

### Dissociation between geometry and arithmetic

Actively listening to geometric statements as opposed to arithmetic sentences selectively activated a single region - the left anterior lateral occipital cortex (aLOC) at the posterior border of the inferior temporal gyrus (figure 3A). Based on adults’ activation maps(*54*), we defined two geometry-responsive ROIs (figure S4C) and compared children’s β-values across the four conditions (geometry, arithmetic, general knowledge, social, figure 3B). Linear mixed-model regressions confirmed that the left aLOC was significantly more activated by geometric than by either arithmetic (t(311.71) = 6.17, p_FDR_ < .001), general knowledge (t(311.71) = 2.63, p_FDR_ = .013), or social statements (t(311.71) = 6.95, p_FDR_ < .001). In contrast, the right aLOC showed significantly higher activation for geometry only when compared to social sentences (t(310.69) = 3.19, p_FDR_ = .004), but not in comparison to arithmetic (t(310.69) = 0.61, p_FDR_ = .54), or general knowledge (t(310.69) = 1.72, p_FDR_ = .10).

**Figure 3.**
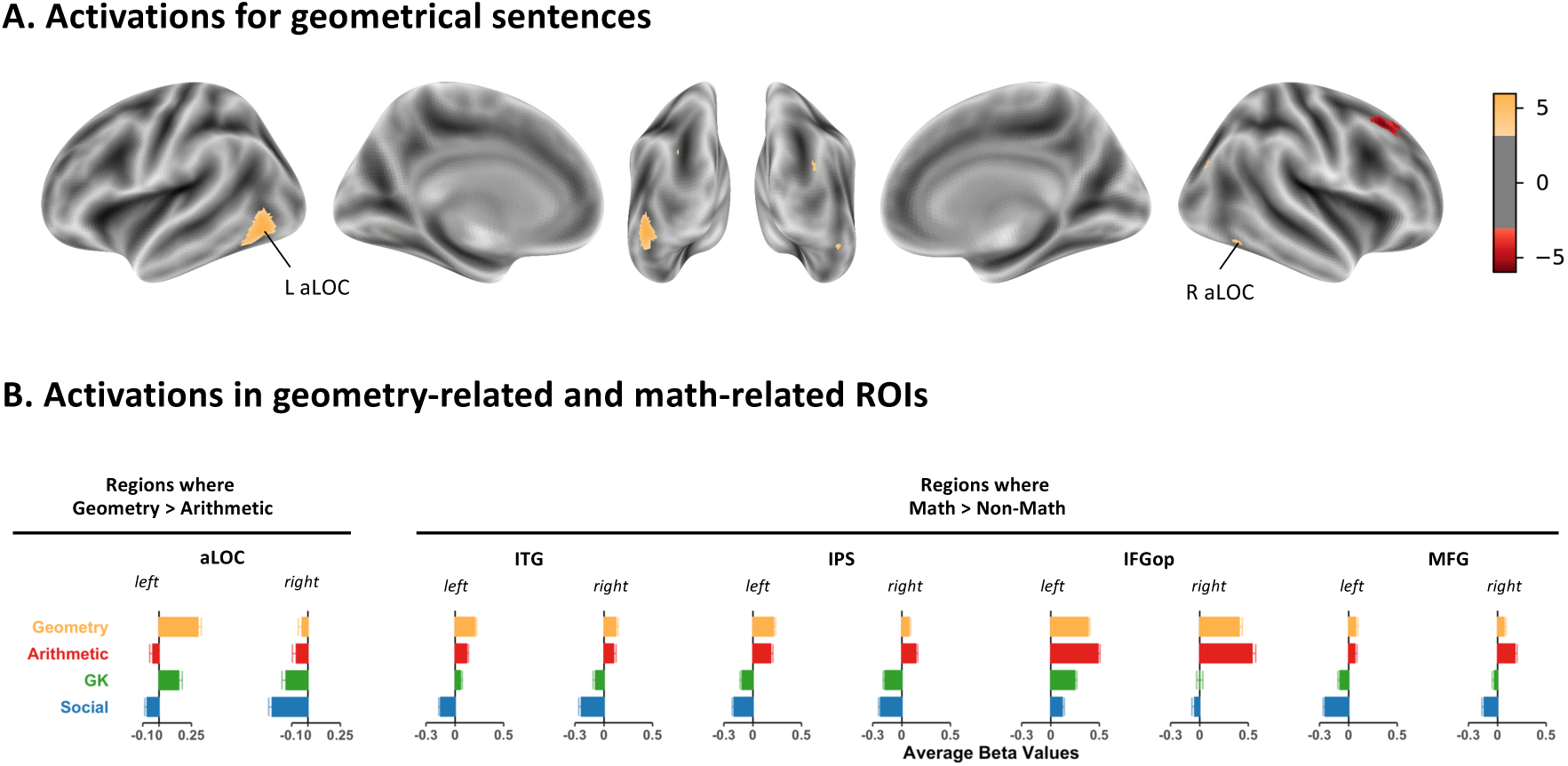
Dissociation between geometry and arithmetic. **(A)** fMRI contrast between geometric and arithmetic sentences across all sessions. Vowel-wise threshold p < 0.001, FDR-corrected α < 0.05. **(B)** Average *β*-values within geometric-related ROIs (left and right aLOC) and in math-related ROIs defined in adults(*54*).

Despite this local difference, arithmetic and geometric responses overlapped in a core common math-responsive network (Figure 3B, figure S7A). Similar mixed-model and Bayesian regressions conducted within the eight math-responsive ROIs revealed no significant differences between geometry and arithmetic after FDR correction (table S3). Moreover, an anterior-posterior dissociation emerged within both IPS, with anterior sections responding more to geometry and posterior ones to arithmetic (figure S7B). Mixed-model regressions with y-coordinates as fixed effect and participant as random effect, confirmed a significant decreased of Δβ-values (*β_Arithmetic_* - *β_Geometry_*) along the y-axis (t = −6.40, p < .001 in the left IPS; t = −4.96, p < .001 in the right IPS).

### Development of the math-responsive network

#### Age-related increases in mathematical selectivity

To track the development of the math-responsive brain network, we examined how mathematical selectivity evolves as a function of age in the eight adult math-responsive ROIs (figure 4A). Mathematical selectivity was defined as the difference (Δ*β*-values) between the activation elicited by math sentences and the mean activation elicited by non-math sentences. Linear mixed-model regression revealed significant age-related increases in math selectivity in three regions: both IPS (left: t(72.06) = 2.52, p = .014, p_FDR_ = .037; right: t(62.80) = 2.83, p = .006, pFDR = .032), and the left IFGop (t(59.45) = 2.73, p = .008, p_FDR_ = .032) (table S4). When hemisphere was entered as an additional factor within regression, no significant Age x hemisphere interaction was observed (all t<1), indicating comparable developmental trajectories across hemispheres. Similar analysis comparing geometry-related activations against all other conditions (arithmetic, GK and social) in left and right aLOC, found no significant relationship with age (t <1 in both aLOC). Finally, control analyses including head motion and tSNR as covariates confirmed that the age-related changed in Δ*β*-values reflected genuine developmental effects rather than differences in data quality (tables S5-S6, SOM).

**Figure 4.**
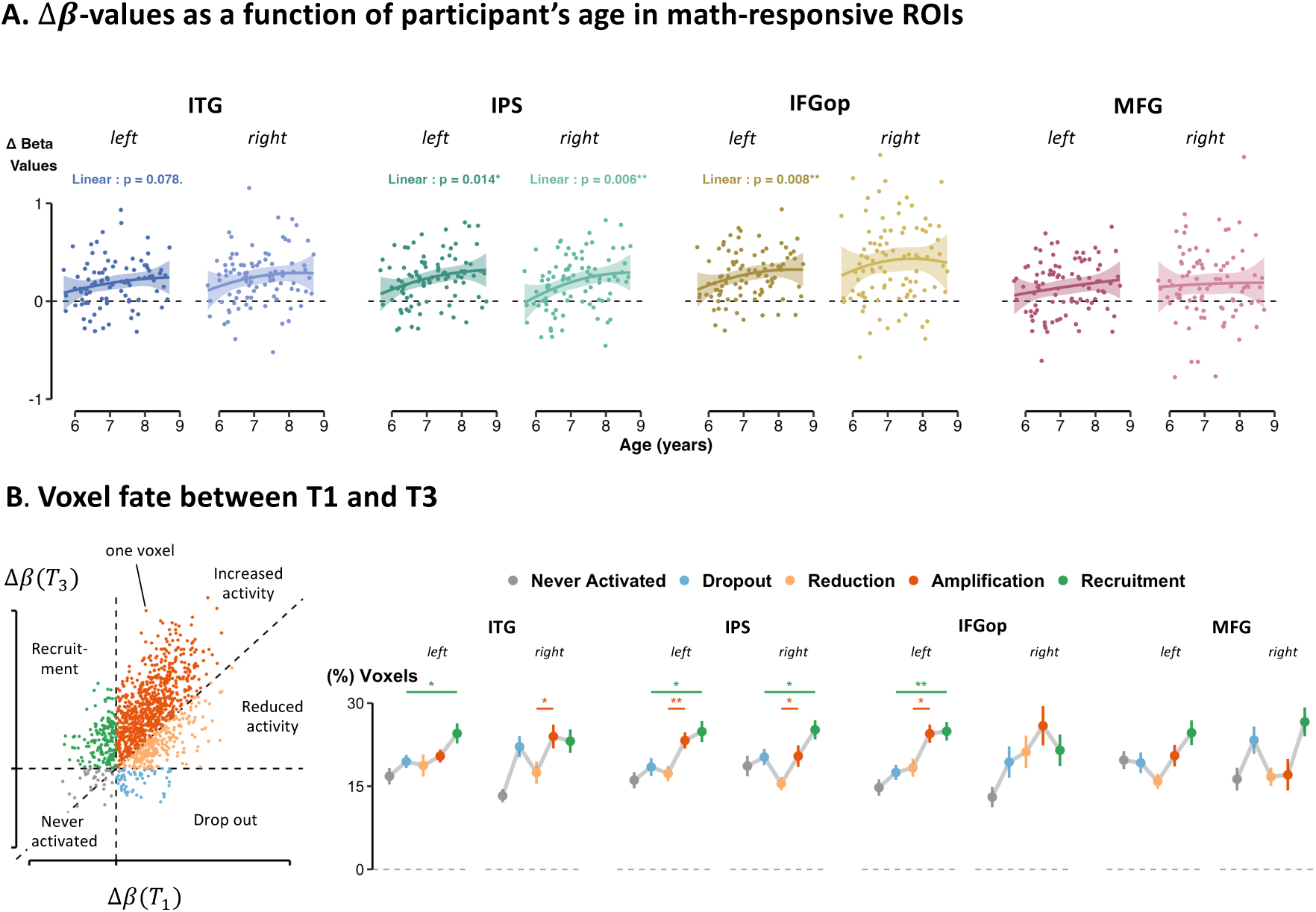
Development of the main areas of the math-responsive network. **(A)** Δ*β*-values (*β_Math_* – *β_Non-Math_*) as a function of children’s age, within each math-responsive ROI. Mixed-model regressions were conducted within each ROI, with age as fixed effect and participant as random effect (*•p < 0.1; *p < 0.05; **p < 0.01; ***p < 0.001*). **(B)** Voxelwise math-specificity changes in children scanned at both T1 and T3 (N = 26). (Left) Model of voxel-activation changes. Each voxel was categorized into five categories based on its activation at T1 and T3: *never activated voxels*, *dropout voxels*, *reduced voxels, amplified voxels,* or finally *recruited voxels*. (Right) For each participant, we computed the percentage of voxels in each of the five categories. We reported the group averages for each of the math-responsive ROI. Linear regressions compared amplified vs. reduced voxels and recruited vs. dropout voxels (*\*p < 0.05; **p < 0.01; ***p < 0.001*).

### Age, schooling or numeracy?

While we used age as a proxy for child development so far, other factors such as schooling (number of days since school entry) and math ability may also shape brain activation patterns. Using similar mixed-model regressions, we observed that mathematical selectivity (Δ*β*-values) was significantly predicted by children’s scores in a numeracy screener test taken prior to each fMRI session, in four of the mathematical ROIs (figure S8A): left IFGop (t(72.08) = 3.06, p = .003, pFDR = .024), both IPS (left: t(85.92) = 2.19, p = .031, p_FDR_ = .062; right: t(73.66) = 2.77, p = .007, p_FDR_ = .028), and left ITG (t(76.36) = 2.34, p = .028, pFDR = .062) (table S7). Interestingly, only one region (left IPS) showed a significant relationship between mathematical selectivity and reading ability (t(67) = 2.97, p = .004, p_FDR_ = .032; figure S8B, table S8).

Note that, unexpectedly, all developmental measures were highly intercorrelated (r=0.92 between schooling and age, r = .82 between age and math ability, and r = .77 between schooling and math ability). To disentangle their unique contributions, we compared mixed-effects models including either schooling, chronological age, or math ability as the sole fixed predictor (participant as a random effect; figure S8C) and estimated Akaike model weights to quantify how well each model explained the Δβ-values. Mathematical ability emerged as the strongest predictor in the left ITG (63%) and left IFGop (59%), whereas in bilateral IPS, model probabilities were more evenly distributed. In the remaining four regions, none of the predictors showed significant effects (all p > .05).

### Changes underlying the increase in math-related activation

What drives the age-related increase in mathematical selectivity observed in certain brain regions - an increase in the maximal activation of the same voxels, or the recruitment of additional cortex (figure 1b)? To investigate this, we tracked the fate of individual voxels across development. We capitalized on the longitudinal aspect of our data and examined the activations of participants who performed both T1 and T3 sessions (N = 26). Within each ROI, we categorized voxels into five categories based on their math-selective activation (Δ*β* = *β_Math_* – *β_Non-Math_*) at T1 and T3 (figure 4B): *never activated voxels* (Δ*β*(*T*_3_) < 0, and Δ*β*(*T*_1_) < 0), *dropout voxels* (Δ*β*(*T*_3_) < 0, and Δ*β*(*T*_1_) > 0), *recruited voxels* (Δ*β*(*T*_1_) > 0, and Δ*β*(*T*_2_) < 0) and finally, within the voxels activated in both periods (Δ*β*(*T*_2_) > 0 and Δ*β*(*T*_1_) > 0), those whose activity was *reduced* (Δ*β*(*T*_1_) < Δ*β*(*T*_2_)) or *amplified* (Δ*β*(*T*_1_) > Δ*β*(*T*_2_)). For each participant, we computed the percentage of voxels in each of these five conditions.

If the math-related signals were stable over development, the distribution of voxels responses in figure 4B should be symmetric around the diagonal Δ*β*(*T*_2_) = Δ*β*(*T*_1_). This null hypothesis was clearly rejected due to both amplification and, to a lesser degree, recruitment. Indeed, the percentages of amplified voxels was significantly higher than the percentage of reduced voxels in the left IFGop (t(25) = 2.62, p = .01, pFDR = .046), both IPS (left: t(25) = 2.92, p = .005, p_FDR_ = .042; right: t(25) = 2.18, p = .034, pFDR = .068) and the right ITG (t(25) = 2.22, p = .031, pFDR = .068) (figure 4B & table S9). We also found modest evidence for recruitment: in left IFGop, left ITG, and bilateral IPS, the percentage of recruited voxels exceeded that of dropout voxels, although these comparisons did not survive FDR correction, except for the left IFGop (t(25) = 3.45, p = .0012, pFDR = .009; table S10). Taken together, these findings suggest that age-related increases in math responses are driven both by enhanced responses within the already-active voxels, and the recruitment of new voxel, although the latter effect seems less pronounced.

### Sentence-level analysis

#### Improved comprehension correlates with decreased activation

While we observed rapid changes in the math-responsive network between ages 5 and 8, it is unlikely that all mathematical statements engage these regions uniformly. More complex sentences may recruit these regions only in older children who possess the necessary conceptual understanding. Conversely, simpler statements may evoke stronger activation in younger children, while in older children, their comprehension may have become sufficiently automated to require fewer neural resources (*automatization* model in figure 1B).

To test these possibilities and examine how changes in brain activation were related to changes in sentence comprehension, we focused our analysis on the sentences that were heard twice by the same children at both T1 and T3. For each sentence of the three domains and each child, we computed the change in brain activation (Δ*β*-values) in both math- and social-related ROIs, by subtracting the *β*-values at T3 from those at T1. For each sentence, we also computed the average behavioral performance across children at both T1 and T3 and used the difference (Δ*Perf*) as a proxy for change in comprehension. Across mathematical sentences, Δ*β*-values, averaged across children and math ROIs, decreased significantly as Δ*Perf* increased (t(19) = −2.79, p = .012, p_FDR_ = .036). This was not the case for general knowledge (t(19) <1) or social sentences (t(21) <1) (figure 5A). This effect was significantly stronger for math compared to social sentences (t(59) = 2.03, p = .047) and marginally greater compared to general knowledge ones (t(59) = 1.72, p = .09). Importantly, when performing the same analysis but within social-related ROIs, no significant relationships emerged (Math: t(19) = 1.45, p = .16, p_FDR_ = .49; GK: t(19) <1; Social: t(21) <1) (figure S9A). This pattern points to a robust relationship between improved comprehension and decreased activation within math-specific regions: the better a mathematical sentence was understood over time, the less activation it elicited in the math-specific regions.

**Figure 5.**
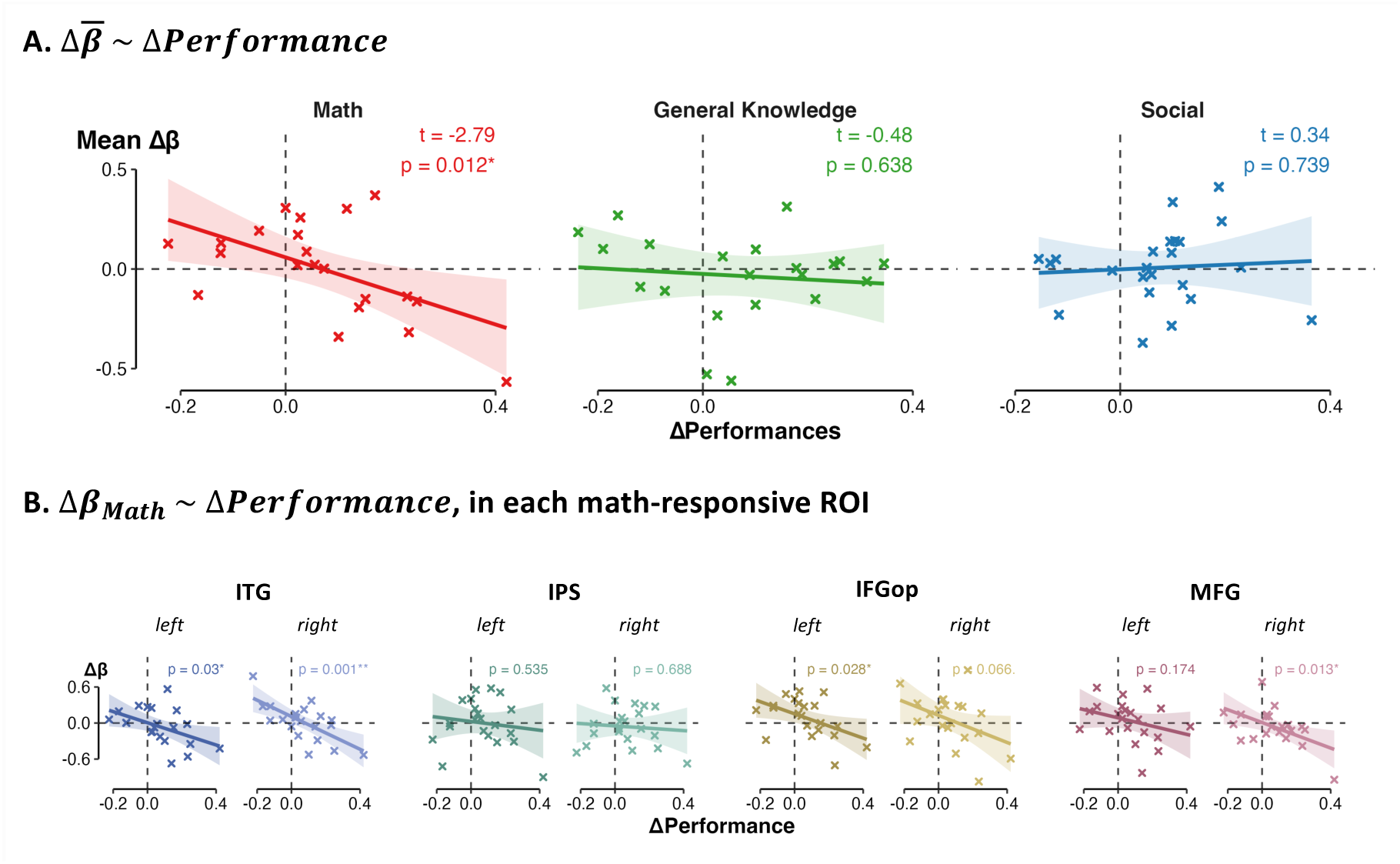
Improved comprehension leads to reduced activation for mathematical sentences in math ROIs. Only children scanned both at T1 and T3 were included here. **(A)** Mean Δ*β*-values (*β*(*T*_3_) − *β*(*T*_1_)) for each condition, averaged across children and math-responsive ROIs, as a function of changes in performance (*\*p < 0.05; **p < 0.01; ***p < 0.001*). **(B)** Mean Δ*β*-values for math sentences and within each ROI, averaged across children, as a function of changes in performance (*\*p < 0.05; **p < 0.01; ***p < 0.001*).

Examining each of the eight math ROIs separately (figure 5B), we observed that all regions, apart from left and right IPS, showed this pattern, with significant decreases in four regions (left and right ITG, left IFGop and right MFG, table S11, although only the right ITG survived FDR correction). In contrast, none of the social regions showed significant decreases in activation–performance relationships for mathematical sentences (all p > .05, except for left TP but with t > 0; table S12, figure S9B).

### An increase in dimensionality in Math Rois

Finally, to go beyond mere activation changes and investigate *how* information is represented at the neural level, we studied developmental change in high-dimensional neural space (figure 1C). We measured their *intrinsic dimensionality* (ID), a quantitative measure of how many independent vector directions in cortical space are required to represent a set of distinct sentences. ID was calculated for each child, condition and each set of ROIs (in both math- and social-related ROIs) (figure 6A). To assess how neural dimensionality evolves with age within each set of ROIs, we performed similar analyses as before but given that each participant contributed multiple measurements across brain regions and age points, the model included random intercepts and slopes for age per participant and a random intercept for ROI.

**Figure 6.**
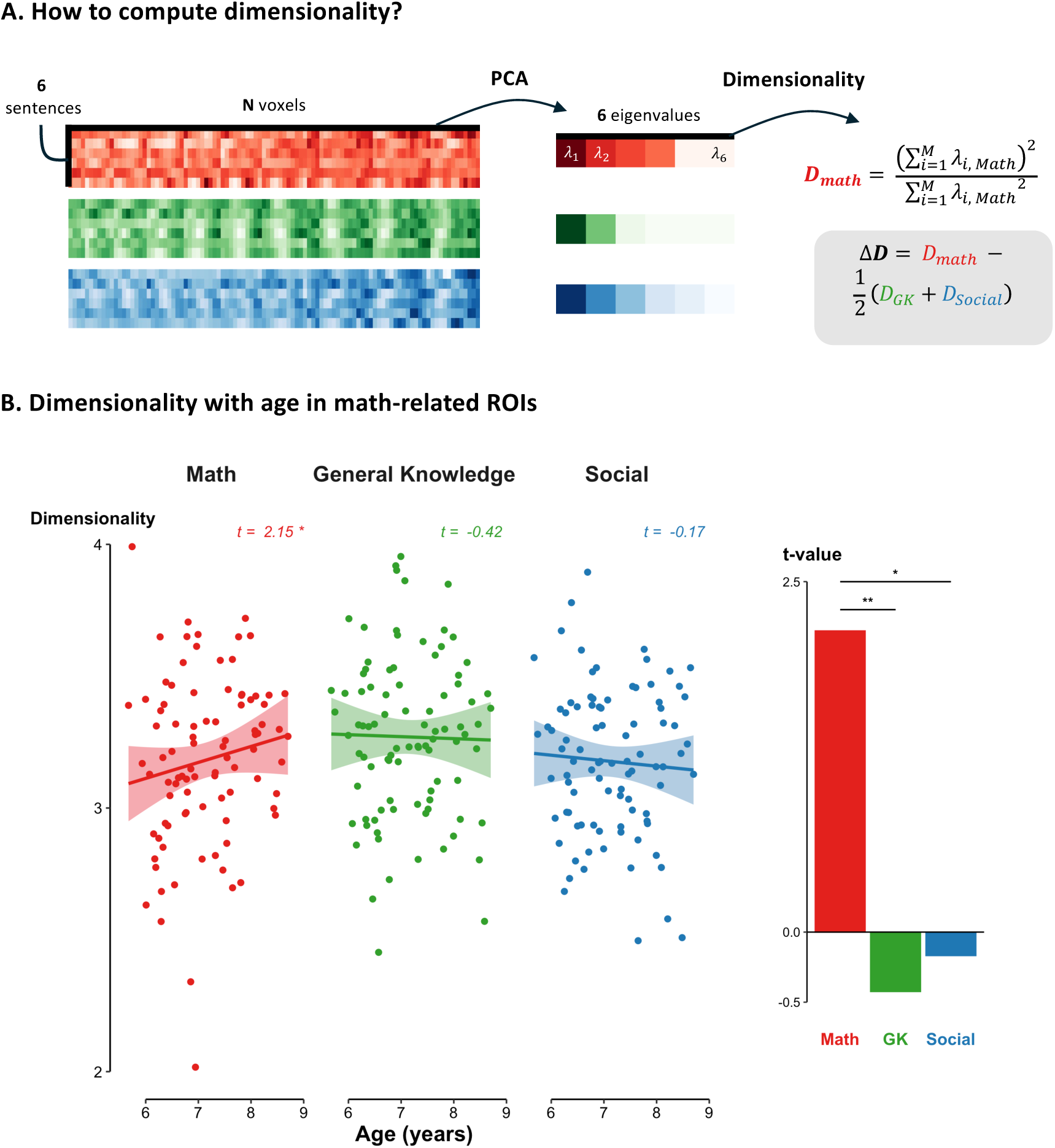
A slight increase in the differentiation of neural space for mathematical concepts. **(A)** For each run performed by each child, for each condition (Math, general knowledge, Social) and within each ROI, we performed a Principal Component Analysis (PCA). Eigenvalues were used to compute the intrinsic dimensionality (ID), reflecting how many dimensions represent the activation patterns. **(B)** Dimensionality values – averaged across ROI - as a function of age, in each condition. Mixed model regressions were conducted, using age as fixed effect, and including random intercepts and slopes for age per participant, and a random intercept for ROI (*\*p < 0.05; **p < 0.01; ***p < 0.001*).

In math-related ROIs, we observed a small but significant age-related increase in dimensionality for math sentences (t(29.67) = 2.15, p = .040), and a non-significant decrease for general knowledge and social sentences (both t <1, figure 6B). In social-related ROIs, no significant age-related changes were observed for any condition (Math: t(29.20) = 1.52, p = .14; GK and Social: t<1) (figure S10A). When condition was added as a predictor, a significant age x condition interaction emerged in math-related ROIs between math and general knowledge sentences (t = −3.03, p = .002) and math and social sentences (t = −2.46, p = .014), but not in social-related ROIs (Math vs. GK: t = −1.25, p = .21; Math vs. Social: t = −1.52, p = .13). These results suggest that, during the first two years of schooling, math-responsive brain regions exploit a higher-dimensional neural space to represent mathematical information, while social-responsive brain regions did not show such developmental change in our age range.

We then performed linear regression in each of the eight math-related ROIs individually (figure S10B). Although most regions showed a positive age-related increase in Δ*Dimensionality* (i.e., differences between math dimensionality and non-math dimensionality), only two – the left ITG and right MFG – showed a statistically significant effect (left ITG: t(89) = 2.20, p = .029, right MFG: t(89) = 2.48, p = .014) which did not survive FDR correction (p_FDR_ = .12 and .11 respectively) (table S13).

## Discussion

We investigated fMRI responses in 6-9 year-old children using a sentence-listening paradigm previously used only in adults (*22*, *24*, *42*). Our findings provide compelling evidence that the neural architecture underlying mathematical cognition is already well-established prior to schooling, including a partial dissociation between arithmetic and geometry. Over the first two years of school, developmental changes include: increased peak activation for math relative to non-math in bilateral IPS, left IFGop and right ITG; an expansion of cortical territory in bilateral IPS, left IFGop and left ITG; activation decreased when sentences meaning became better mastered, in bilateral ITG, left IFGop and right MFG; and a slight increase in the dimensionality of neural space encoding math sentences (figure 7). We now discuss these points in turn.

**Figure 7.**
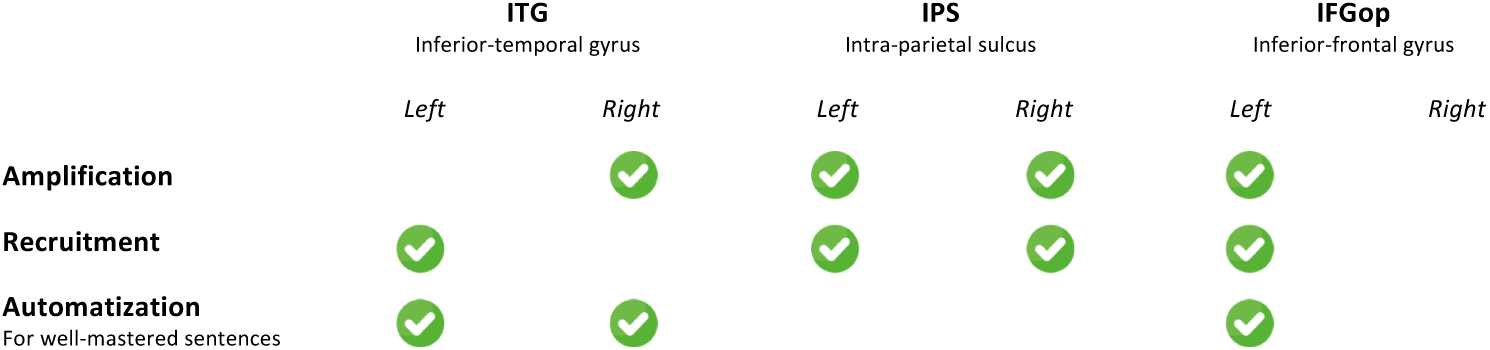
Summary of developmental changes in cortical space. Green crosses indicate significant changes.

Our study extends to young children, including preschoolers, the existence of a network of brain regions selectively engaged when actively processing mathematical sentences compared to non-math ones (*22*, *42*). It includes a well-documented fronto-parietal network but also extends to regions less frequently emphasized in the literature, including the bilateral inferior temporal gyrus and bilateral posterior cingulate (figure 2). This early mathematical brain network closely mirrors the well-characterized math-responsive network in adults (*15*, *16*, *24*, *25*) and children (*35*, *47*, *69*). The present findings indicate that this network can already be engaged, not only by concrete mathematical objects such as sets of dots, but also by purely verbal symbols. Thus, even at the very beginning of schooling, mathematical language is not only understood but also functionally integrated and capable of recruiting a math-responsive cortical network. In agreement with the triple-code model of number processing (*70*), bilateral IPS appears as a semantic convergence zone capable of activating from purely verbal inputs, and now seen to do so (1) since preschool (2) for both arithmetic and geometry, and (3) in tight coordination with bilateral ITG, IFG and MFG.

While several studies have probed the neural bases of symbolic vs. non-symbolic numerical cognition (*32*, *33*, *35*, *69*) relatively few directly compared different mathematical subfields (*71*). A recent study showed broad overlap in brain activation when children compared numbers, geometric shapes, or performed a number line task (*47*). In adults, however, while there is broad overlap in brain activations to statements on geometry, algebra, trigonometry or complex numbers (*22*), the anterior LOC responded selectively to geometric sentences. This region has since been shown to be implicated in other visuospatial mathematical tasks, such as graph interpretation (*72*) and geometric shape processing (*58*). Here, our study extends these observations to children with the left aLOC showing greater activation for geometrical sentences as opposed to arithmetic ones (figure 3). Moreover, we showed that the left aLOC’s selectivity for geometry remained stable across development, with no indication that the right aLOC would develop similar geometry-specific responses over time. Another study, however, showed greater activation in the right aLOC in adults than in 6-year-olds during the actual perception of geometric shapes (*58*) and a positive correlation between right aLOC activity evoked by visual graphics and math education (*73*). Together, these results suggest that the left aLOC may show a greater involvement in acquiring geometry-related words and sentences, while the right aLOC would accumulate knowledge of visual shapes and graphics – a hypothesis that should be tested in future work.

Our results also shed light on the complex issue of the developmental dynamics of prefrontal engagement. While some studies report a developmental increase in prefrontal activation (*35*, *37*, *50*, *51*), others argue that mathematical cognition progressively shifts away from frontal regions toward greater parietal reliance (*27*, *30*, *32*, *33*, *36*, *47*, *74*). By tracking the same children over two years, we observed both an age-related increase in left IFG activation (figure 4; alongside bilateral IPS increases), but also a decrease in bilateral prefrontal activation for sentences that became better understood from T1 to T3 (figure 5; while IPS activity remained stable). Rather than being contradictory, these effects reflect distinct phases of a dynamic developmental trajectory. Prior studies documenting a decline in frontal activity with age have primarily relied on cross-sectional comparisons between children and adults, thus capturing only the endpoints of the developmental continuum. In contrast, studies reporting an age-related increase in frontal engagement, like the present one, have typically compared children across different ages, offering a finer-grained perspective on the developmental dynamics. Overall, the data are compatible with a quadratic model in which frontal engagement initially increases during the acquisition of formal mathematical abilities, then gradually declines as mathematical processing become automatized (*51*). Similar trajectories have been observed in the hippocampus during arithmetic processing (*53*), and in parieto-frontal networks during reading acquisition (*63*), suggesting a broader principle whereby an initial frontal engagement facilitates the acquisition of complex skills before neural reorganization optimizes processing efficiency and activation shifts towards more specialized regions as expertise emerges.

Since the bilateral IPS was the only region showing both an age-related increase in math activation (figure 4) and no significant decrease for well-mastered sentences (figure 5), our findings confirm its central role at the core of mathematical cognition (*1*, *4*, *15*, *16*, *25–27*, *75*) (figure 7). We found no evidence of hemispheric asymmetry, consistent with recent findings (*35*, *43*, *45*). Whereas prior evidence has highlighted the left IPS as a key site for acquiring symbolic mathematics (*30*, *32*, *37–41*, *74*), evidence for a parallel development in the right IPS has been less consistent (*33*, *36*). By investigating semantic processing of mathematical sentences, our study suggest that even though the right IPS is already engaged in numerical cognition from infancy (*1*, *26*), it continues to develop alongside the left IPS during the acquisition of formal mathematical reasoning.

We found that not only age, but also school exposure and mathematical ability significantly predicted increased math-specific activity in key math-selective regions, including the left ITG, left and right IPS, and left IFGop. These results mirror previous studies reporting modulations of math-induced brain activity by mathematical ability (*37*, *39*, *43*, *45*, *76*). Using model comparisons, we attempted to disentangle these developmental components and showed that mathematical ability was the strongest predictor in the left ITG and IFGop, whereas in the IPS, no single predictor clearly dominated. Given the strong intercorrelations among these developmental variables, isolating their distinct effects remains challenging. Future work will be needed to fully address this issue by scanning populations in whom age and math level are dissociated, as was done in the field of reading acquisition (*21*, *77*).

Our data also clarify competing theories of math development and provide a more nuanced view of how the math-responsive brain network evolves during early schooling (figure 1). Across the first two years of schooling, we indeed observed all three forms of developmental change illustrated in figure 1B: age-related *amplification* of activation in bilateral IPS, left IFGop and right ITG (figure 4B); modest *recruitment* of additional voxels in largely the same regions (figure 4B); and reduced activation for well-mastered facts in bilateral ITG, left IFGop, right MFG but notably not in IPS (figure 5B). These mechanisms are thus not mutually exclusive but characterize complementary aspects of developmental change. The recruitment of new cortical territory for mathematics raises important questions for future research. What was the function of these voxels before becoming tuned to mathematical content? Were they unresponsive, domain-general, or selectively engaged in other functions, only to be later repurposed for math processing, in line with the neuronal recycling hypothesis (*21*)? This question remains largely unexplored in the field of mathematical cognition, in contrast to reading research (*63*, *78*, *79*). One plausible hypothesis is that IPS sites initially involved in visuo-spatial operations are repurposed for encoding numerical or geometric content – a possibility supported by the known overlap of these representations in the IPS of both children and adults (*47*, *71*, *80*, *81*).

Finally, our sentence-based design allowed us to probe developmental change in neural space (figure 1C). Inspired by advances in the theoretical and empirical description of coding by high-dimensional neural subspaces (*64–67*), we estimated the intrinsic dimensionality of the neural space occupied by the same mathematical sentences over time. We found a small yet significant increase in dimensionality in math-related regions (figure 6). This finding suggests that, as children mature, the complexity of the neural manifold that encodes mathematical contents increases. This finding offers a potential solution to the vexing question of how new mathematical facts, including advanced ones, can be acquired within the same overall cortical circuit: new concepts can be encoded using previously unused directions of neural vector space. This interpretation remains tentative, however, as the increase in dimensionality did not reach significance within specific math-related ROIs after FDR correction. Replication with higher signal-to-noise methods will therefore be essential, using either ultra-high-field imaging or human intracranial recordings, which have recently been shown to capture various semantic dimensions of words and sentences (*82*, *83*), including numbers (*84*).

To our knowledge, no previous study has examined how spatial neural complexity evolves during development—except for one, which used representational similarity analysis and found less differentiated neural representations in children with math learning difficulties compared to typically developing peers (*76*). Temporal complexity has been more extensively explored, especially through entropy measures in electrophysiology (*85–87*) and fMRI (*49*, *88*). These studies generally report increasing temporal complexity from childhood to adulthood, suggesting that the time domain may offer an additional axis for coding flexibility during development.

The present methods, involving longitudinal recordings of brain activity patterns evoked by the same sentences over development, offer a new tool for conceptualizing and measuring the impact of education on the brain. While we focused here on mathematical development, this dataset also holds rich potential for exploring developmental changes in social, linguistic, and musical cognition.

## Materials and Methods

### Participants & Task

#### Ethics

All experiments were approved on December 21, 2022, by the French national ethical committee (CPP: *Comité de Protection des Personnes,* CPPIDF-2022-MS276).

#### Participants

58 children were recruited in spring 2022 and participated in an initial MRI session (***T1***) during summer-fall 2022. At the time, children were 5-6 years old, either finishing kindergarten or entering first grade. Five were excluded after T1: one did not wish to continue, one had attention disorders, and three had language disorders (exclusion criteria). 53 participants (27 boys, 26 girls, mean age = 6.38 ± 0.35 years, ages range = [5.67, 6.99]) were therefore included in T1 analysis. One year later (May–October 2023), 46 of these participants (24 boys, 22 girls, mean age = 7.16 ± 0.30 years, ages range = [6.57, 7.60]) returned for a second session (**T2**). Finally, 35 of them (16 boys, 19 girls, mean age = 8.07 ± 0.33 years, ages range = [7.47, 8.70]) returned for a third session (***T3***), from May to October 2024, corresponding to the end of second grade or start of third grade. Participants were recruited from schools in the priority education network in the north of Île-de-France (serving low-income families). On average, mothers had 0.92 years of education beyond high school, with 25 on 53 having a *baccalauréat* (BAC) or less, while fathers had 0.28 years beyond high school, with 31 on 53 having a BAC or less. Sample sizes were not predetermined using statistical methods, but were comparable to or larger than those used in recent cross-sectional or longitudinal studies (*35*, *38*, *47*, *63*, *79*).

#### Stimuli

72 sentences (24 in each cognitive domain) and 24 melodies were created. The sentences covered three cognitive domains: mathematics (e.g., “Three plus three is six”; hereafter called Math), general non-mathematical knowledge about biology (e.g., “The ant is a black insect”; called General Knowledge), and social sentences (e.g., “According to the policeman, thieves are nice”, called Social). Within each domain, 12 statements were true and 12 were semantically false. All statements were recorded by a female native French speaker and were matched for syntactic structure, number of words (in average 7.5 words per sentence for both math, general knowledge and social), number of phonemes (in average 19.9, 20.2 and 22.6 phonemes per sentence for respectively math, general knowledge and social), duration (in average 3.46s, 3.52s and 3.46s for respectively Maths, general knowledge, Social), and mean word frequency in French. A detailed list of statements is presented in supplementary figure S1A.

The melodies were equalized in duration and intensity, and half of the melodies were consonant. In dissonant melodies, one of the final two notes was shifted one tone away from the expected key. The musical condition is not analyzed in the present article.

#### Paradigm

Children were presented with the sentences and musical melodies in random order and asked to judge whether the sentences were true or false, and the melodies normal or weird. Each trial began with a beep, followed by the presentation of a sentence or melody. At the end of the presentation, participants had three seconds to press a right button for *correct/normal*, or a left button for *incorrect/weird*. Visual and auditory feedback were provided at the end of the three seconds, followed by an inter-trial interval varying between 1 and 3s (average = 2secs). The whole experiment included 4 runs of 24 pseudo-randomized trials (8 sentences/melodies, with 4 correct and 4 incorrect, from each condition), with each run lasting approximately 3’30”. However, as children could opt out, they almost never completed the entire experiment: 1.54 ± 0.30 runs/child at T1; 1.85 ± 0.33 at T2; and 2.29 ± 0.40 at T3.

#### Data quality and inclusion criteria

A total of 30% of T1 runs, 46% of T2 runs, and 27% of T3 runs were excluded from the analysis due to excessive motion (i.e., > 3mm). The final sample included 40 children in T1, 24 in T2 and 30 in T3, each with at least one acceptable run, for a total of 94 time points. Among the included children, the overall amount of motion (averaged across axes and runs) did not significantly differ across periods (T1 vs. T2: t(62.1) = −1.71, p = .21; T1 vs. T3: t(57.4) <1; T2 vs. T3: t(60.3) <1; obtained from linear mixed-model regressions, with participant as random effect), nor were significantly correlated with children’s age (t(66.14) <1), or with the number of days the child spent at school (t(59.55) <1) (figure S2A). Despite this absence of differences, motion parameters were included as regressors in the fMRI data analysis. Motion curves for each subject and each run are reported in the supplementary data. Similarly, the average temporal signal-to-noise (tSNR) – i.e., the mean of the tSNR across voxels - computed for each included scanning did not significantly differ across periods (T1 vs. T2: t(59.7) = 1.41, p = .34; T1 vs. T3: t(55.6) <1; T2 vs. T3: t(57.8) = −1.42, p = .34) and was not significantly correlated with children’s age (t(61.33) <1) or with the number of days the child spent at school (t(55.34) <1) (figure S2B).

#### Experimental procedure

Before each fMRI session, to enhance the quality of data, children were trained in a mock scanner. They listened to a recording of the noises associated with all fMRI sequences and were trained on the fMRI task with 4 example trials. They received live feedback on their motion. Following this ∼20 min practice session, children proceeded to the actual MR scanner. Each session began with the acquisition of anatomical images (see below), followed by one functional run of a visual categories localizer task (not analyzed here), one run of the sentence comprehension task, a diffusion tensor imaging sequence (not analyzed here), and then alternating runs of visual localizer and sentence comprehension tasks until the child opted to end the session. To minimize head motion, the quality of each sequence and functional run was reviewed immediately after acquisition, and children received verbal feedback accordingly.

On the same day as the MRI session, we performed behavioral tests and experiments with the children, to (1) verify that they had no learning disabilities (exclusion criteria), and (2) quantify their individual abilities in different domains (e.g., motricity and oral comprehension). Here, we analyzed two of them, which we describe in the next sections.

### Behavioral Experiments

#### Numeracy Screener

Each child’s mathematical ability was assessed by a behavioral test, the numeracy screener (*89*), conducted on the same day as each of their fMRI session. This paper-and-pencil test evaluates children’s ability to compare both symbolic and non-symbolic numerical magnitudes. During each trial, children were instructed to cross out the larger of two single-digit numerical magnitudes. The test comprised two conditions: a symbolic condition, in which children compared 56 pairs of Arabic numerals, and a non-symbolic condition, where they compared 56 pairs of dot arrays. Each condition lasted two minutes, with the order of conditions randomized across participants. For each condition, the number of errors was recorded, along with the time taken to complete the task (if completed within the two-minutes limit). An overall mathematical ability was then computed for each child, at each time point, as:

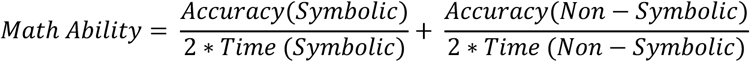

#### Reading fluency

Reading fluency was quantified as the mean number of correctly read words during a one-minute timed reading task (*90*).

### fMRI parameters and analysis

#### MRI Data Acquisition

MRI acquisition was performed on a 3-Tesla scanner (Siemens Prisma) equipped with a 64-channel head-coil. T1-weighted structural images were acquired (repetition time (RT) = 2300ms; echo time (TE) = 2.98ms; voxel-size = 1mm^3^). Functional images were obtained using a high-resolution multiband imaging sequence (RT = 1810ms; TE= 30.4ms; voxel-size = 2mm^3^; multiband factor = 3). Each run comprised 142 functional scans covering the whole brain (69 slices). To estimate distortions, two spin-echo field maps with opposite phase encoding directions were acquired: one volume in the anterior-to-posterior direction (AP) and one volume in the other direction (PA). Children were wearing noise-protection earphones.

#### MRI preprocessing

Preprocessing was performed with the standard pipeline fMRIprep (version 22.0.2). The following description was generated by fMRIprep.

Anatomical Data: The T1-weighted (T1w) images was corrected for intensity-non-uniformity (INU) and used as T1w-reference throughout the workflow. The T1w-reference was then skull-stripped using a target template (OASIS30ANTs). Brain tissue segmentation of cerebrospinal fluid (CSF), white-matter (WM) and gray matter (GM) was performed on the brain extracted T1w. Volume-based spatial normalization to two standard spaces (MNI152NLin6Asym, MNI152NLin2009-cAsym) was performed through nonlinear registration using brain-extracted versions of both T1w reference and the T1w template.

Functional data preprocessing: First, a reference volume and its skull-stripped version were generated using a custom methodology of fMRIPrep. Head-motion parameters with respect to the BOLD reference (transformation matrices, and six corresponding rotation and translation parameters) were estimated before any spatiotemporal filtering. BOLD runs were slice-time corrected. The BOLD time-series (including slice-timing correction) were resampled onto their original, native space by applying the transforms to correct the head-motion. The BOLD reference was then coregistered to the T1w reference. Co-registration was configured with six degrees of freedom. Several confounding time-series were calculated: framewise displacement (FD), DVARS (Derivative Variance Across Space) and three region-wise global signals. FD and DVARS are calculated for each functional run. The three global signals are extracted within the CSF, the WM, and the whole-brain masks. These steps were performed for each of the BOLD runs.

#### fMRI GLM Models

fMRI first-level models were estimated for each subject to model task-related blood-oxygen-level dependent (BOLD) responses. The design matrix included four experimental conditions: math, general knowledge, social, and music. Two regressors were included to account for button responses (left, right). Confound regressions included six motion corrections to account for head translation and rotation along the three axes, as well as three regressors for physiological noise: average signal across all brain voxels, average CSF signal, and average WM signal. To account for low-frequency signal drifts, polynomial drift models from constant to 5^th^ order were added. Finally, a constant term was also included to model the baseline signal.

Group-level analysis was performed using a second-level model with spatial smoothing applied at a full-width half-maximum (FWMHM) of 8mm. Unless indicated, all brain activation results are reported using a voxel-wise threshold of p < 0.001, corrected for multiple comparisons across the whole brain using false discovery rate (FDR) at α < 0.05.

We performed two additional control analyses to ensure that the math-responsive network was independent of sentence complexity: 1) we repeated the first-level analysis using only trials that were correctly judged by children (figure S3A), and 2) we added a trial-by-trial RT-difficulty regressor, computed as each trial’s RT minus the participant’s mean RT, to the first-level model (figure S3B).

#### Regions-of-Interests from adults’ fMRI data

To assess whether the children’ neural networks resemble those observed in adults, we used fMRI data collected in a cohort of adult participants, with a similar paradigm (*54*). In this paradigm, adults were asked to evaluate the truthfulness of sentences belonging to different semantic conditions (e.g., mathematics). From the activation maps obtained for the contrast math vs. non-math sentences at the group-level, we defined eight *math-responsive* regions-of-interest (ROIs): left and right intraparietal sulcus (*IPS*), left and right inferior temporal gyrus (*ITG*), left and right inferior frontal gyrus *pars opercularis* (*IFGop*) and left and right medial frontal gyrus (*MFG*) (figure S4A). Similarly, from the activation maps obtained for the contrast social vs. contextual knowledge sentences (i.e., referring to a person vs. facts in a specific context: “*For Einstein the atomic bomb is a harmless object”* vs. “*In the Amazon spiders are harmless animals”*), we defined seven *social-responsive* regions-of-interest: bilateral temporal gyrus (TP), left angular gyrus (L AG), right temporo-parietal junction (R TPJ), left medial frontal gyrus (L MFG), medial frontal gyrus (MedFG) and precuneus (figure S4B). Finally, from the activation maps obtained for geometry vs. other mathematical sentences (e.g., “*A rectangle has only one axis of symmetry”* vs. “*Three multiplied by twelve is thirty-six*”), we defined two *geometrical-responsive* regions-of-interest: the left and right anterior lateral occipital cortex (aLOC) (figure S4C). Within each region-of-interest, we identified the 10% of voxels with highest activation peak for adults for the contrast in question (e.g., math vs. non-math). The *β*-values extracted in children in each region was computed from these 10%-voxels.

Children’s fMRI data were normalized to the MNI152NLin2009cAsym template, while the adult data were normalized to the Montreal Neurological Institute avg152 template. To extract the *β*-values from the correct voxels, we transformed the children’s statistic maps into the adult template first, before applying spatial filtering.

### Data analysis

#### Mixed-model regressions

We employed mixed-model regression to account for the hierarchical structure of the data (repeated measures within each participant), and because the number of observations and participants’ ages varied. Mixed-model regressions were used both to 1) characterize the math-responsive network, by examining differences in activations between different conditions (e.g., math vs. social), and 2) investigate the development of this math-responsive network, by studying the evolution of these activations with age.

Degrees of freedom for fixed effects in the linear mixed models were approximated using Satterthwaite’s method, which accounts for both the hierarchical structure of the data and the estimation uncertainty of random effects. These degrees of freedom were then used to estimate p-values for the fixed effects. False Discovery Rate (FDR) correction was applied to control for multiple comparisons using the Benjamini-Hochberg procedure.

All regressions performed are described just below.

To examine differences in performance during development, we conducted a binomial mixed-model regression with trial-by-trial performance as dependent variable, condition (math, general knowledge, social) and age as fixed effects, and participant as a random effect (table S1). Similarly, to examine differences in reaction times during development, we performed a linear mixed-model regression with reaction times in correct trials as dependent variable, condition (math, general knowledge, social) and age as fixed effects, and participants as a random effect (figure S1B, table S2).

To compare brain activations within each of the math-selective ROI, we performed a linear mixed-model regression, with bold signal (%) at each time step as dependent variable, condition (math, general knowledge, social) as fixed effect and participant as a random effect (figure 2B). Similar analyses were conducted separately in each of the periods (T1, T2, and T3; figure S6).

To compare brain activations within each of the geometrical-selective ROI, we performed linear mixed-model regression, with *β*-values as dependent variable, condition (geometry, arithmetic, general knowledge and social) as fixed effect, and participant as a random effect (figure 3B). We conducted similar frequentist linear mixed-model regression, as well as Bayesian ones in the math-selective ROI, comparing only the geometrical and arithmetical *β*-values (figure 3B, table S3). To investigate the brain activations along the y-axis with both left and right IPS, we performed mixed-model regressions with Δ*β*-values (*β_Arithmetic_* – *β_Geometry_*) as the dependent variable, y-axis as the fixed effect and participant as a random effect (figure S7B).

To track the development of math-responsive brain network, we conducted, within each of the math-responsive ROI, a linear mixed-model regression with Δ*β*-values (*β_Math_* – *β_Non-Math_*) as dependent variable, age as fixed effect and participant as a random effect. To capture potential non-linear effects, the model included a second-order polynomial term for age (figure 4A, table S4). Given that the quadratic term was not significant in any of the regions tested, only a linear term was included in subsequent analyses. To evaluate hemispheric differences, we performed similar linear mixed-model regressions with Δ*β*-values (*β_Math_* – *β_Non-Math_*) as dependent variable, age and hemisphere as fixed effects and participant as a random effect, within each set of bilateral math-responsive ROI (e.g., IPS).

Following the approach in Nord et al. (2021), to ensure that these age-related increases in Δ*β*-values were not driven by age-related differences in data quality, we conducted two additional linear mixed-model regressions within each math-selective ROI, including either age and average head motion during, or age and averaged temporal signal-to-noise as fixed effects, Δ*β*-values (*β_Math_* – *β_Non-Math_*) as dependent variable, and participant as random effect (table S5, S6).

To examine other developmental factors, we also conducted linear mixed-model regressions within each math-responsive ROI, with the same dependent variable as above (i.e. Δ*β*-values (*β_Math_* – *β_Non-Math_*), participant as a random effect, and either math abilities or reading abilities as fixed effect (table S7 and S8).

Finally to assess how neural dimensionality evolves with age, we performed, within each set of ROIs (either math-responsive ROIs, or social responsive ROIs) and for each condition (math, general knowledge, and social), linear mixed-model regressions, with dimensionality-values within each ROI as dependent variable, age as fixed effect, and included random intercepts and slopes for age per participant and a random intercept for ROI, to take into account that each participant contributed multiple measurements across brain regions and age points (figures 6B and S10A). We also performed similar regressions, in each of the set of ROIS, with same dependent variables, random intercepts and slopes, but with age and condition as fixed effect. We also performed simple linear regression in each of the math-responsive ROI, without any random effect due to insufficient variance across participants (figure S10B, table S13).

#### Longitudinal analyses

In two analyses, we took advantage of the longitudinal design of our dataset by focusing on participants who completed T1 and T3 sessions (26 children in total). Instead of using mixed-model regressions, we directly examined changes in activation patterns either at the voxel level or for individual sentences. The first analysis tracked voxel-level activation changes within individuals over the two-year interval (figure 4B). Within each math-responsive ROI, voxels were categorized into five categories based on their math-selective activation (Δ*β* = *β_Math_* − *β_Non-Math_*) at T1 and T3: *never activated voxels* (Δ*β*(*T*_3_) < 0, and Δ*β*(*T*_1_) < 0), *dropout voxels* (Δ*β*(*T*_3_) < 0, and Δ*β*(*T*_1_) > 0), *recruited voxels* (Δ*β*(*T*_3_) > 0, and Δ*β*(*T*_1_) < 0) and finally, within the voxels activated in both periods (Δ*β*(*T*_3_) > 0 and Δ*β*(*T*_1_) > 0), those whose activity was *reduced* (Δ*β*(*T*_3_) < Δ*β*(*T*_1_)) or *amplified* (Δ*β*(*T*_3_) > Δ*β*(*T*_3_)). In each participant, we computed the percentage of voxels in each of these five conditions, and we computed simple linear regressions to compare the percentages of *amplified* and *reduced* voxels (table S9), and the percentages of *recruited* and *dropout* voxels (table S10).

The second analysis assessed how brain responses to the same sentences evolved across time in the same children. Each of these 26 children heard an average of 10 ± 2.65 sentences repeated across the two sessions. For each sentence, we computed the average behavioral performance across children at both T1 and T3 and used the difference (Δ*Perf*) as an index of comprehension improvement. For each sentence and each child, we computed the change in brain activation (Δ*β*-values) by subtracting the *β*-values at T3 from those at T1, within each set of ROIs (either math-responsive ROIs or social-responsive ROIs). We then performed simple linear regressions between Δ*β*-values, averaged across children and set of ROIs, and Δ*Perf*, within math-responsive ROIs (figure 5A) and within social-responsive ROIs (figure S9A). We further examined whether this relationship differed significantly across conditions, by including condition as an additional predictor in the model (Δ*β*-values ∼ Δ*Perf* × Condition) and examining the interaction term. We also conducted similar analysis within each math-responsive ROI (figure 5B, table S11) and within each social-responsive ROI (figure S9B, table S12), with Δ*β*-values averaged across children.

We chose to compare T1 and T3, rather than T1–T2 or T2–T3, for two main reasons: (1) these timepoints represent the two endpoints of the longitudinal span, offering maximal contrast, and (2) the sample size of included children was smallest at T2.

#### Comparison of models

We assessed model fit using the Akaike information criterion (AIC), which balances goodness of fit against model complexity (i.e., the number of predictors). The model with the lowest AIC value was considered the best-fitting model. To evaluate the robustness of model differences, we computed the Akaike weights, which estimates the probability that a given model is the best model, considering the evidence from all other models. For each model, the Akaike weight was computed as:

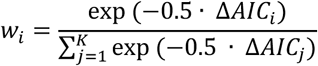

were Δ*AIC_i_* = *AIC_i_* − min (*AIC*).

#### Intrinsic dimensionality

For each run performed by each child, and for each condition (Math, general knowledge, Social) within each region-of-interest or set of region-of-interest, we performed a Principal Component Analysis (PCA; figure 6A). Specifically, we conducted the neural covariance matrix based on a data matrix **B** of shape 6 × *N_Voxels_*, where each row corresponds to one of the six sentences presented during the run, and each column represents a voxel’s activation. When performed within each region-of-interest, N is the number of voxels within this region. When performed within a set of regions, N is the number of voxels within all mathematical or social regions. We then extracted the eigenvalues (*λ*_1_, *λ*_2_, …, *λ*_6_) of this covariance matrix, which indicate the variance explained by each principal component. To estimate the intrinsic dimensionality of the neural activation patterns, we used the “effective dimensionality” or “participation ratio”, defined as

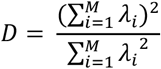

This measure captures how many dimensions are effectively needed to represent most of the data. It reaches its minimum value (1) when all variance is concentrated in a single component (e.g., *λ*_1_ = 1, *λ_i_*_<2_ = 0), and its maximum value (M) when all components contribute equally (i.e., *λ_i_* = *λ_k_*, ∀*∀*).

## Code availability

All code used for data preprocessing, statistical analyses, and figure generation is available on the Open Science Framework (OSF) at https://osf.io/5kvd9/overview?view_only=52f336a5f13f446eb569ada6c798cf31.

## Acknowledgements

This research was funded by the Vareille Foundation, INSERM, CEA, Collège de France, France 2030 program (ANR-23-IAHU-0010) and an ERC grant MathBrain (ERC-2022-ADG101095866) to SD. Neuroimaging experiments and TM’s PhD were paid by the Vareille Foundation. The foundation had no role in the design of the scientific study, data collection, analysis and interpretation, or manuscript preparation. We are sincerely grateful to the NeuroSpin support teams for help in data acquisition, especially the nurses and the fMRI technicians. We also thank all children who participated.

## Supplementary text

### Age-related increases in mathematical selectivity

Linear mixed-model regression revealed significant age-related increases in math selectivity in three of the math-related ROIs: both IPS and the left IFGop. To ensure that these age-related increases in Δ*β*-values were not driven by age-related differences in data quality, we conducted two additional linear mixed-model regressions within each ROI, including either a head motion or temporal signal-to-noise (tSNR), as covariates. Head motion was not significant in any ROI (all p > .12) and age remained a significant predictor in the same regions with nearly unchanged p-values (table S5). Although tSNR emerged as a predictor in two regions – the right ITG (t(83.49) = 2.85, p = .006, p_FDR_ = .051) and the left MFG (t(90) = 2.19, p = .031, p_FDR_ = .099) - age effects persisted in all initially identified regions, again with nearly unchanged p-values (table S6). These control analyses confirmed that Δ*β*-values changes reflect genuine development effects rather than variations in measurement quality over time.

## Supplementary Figures

**Fig. S1.**
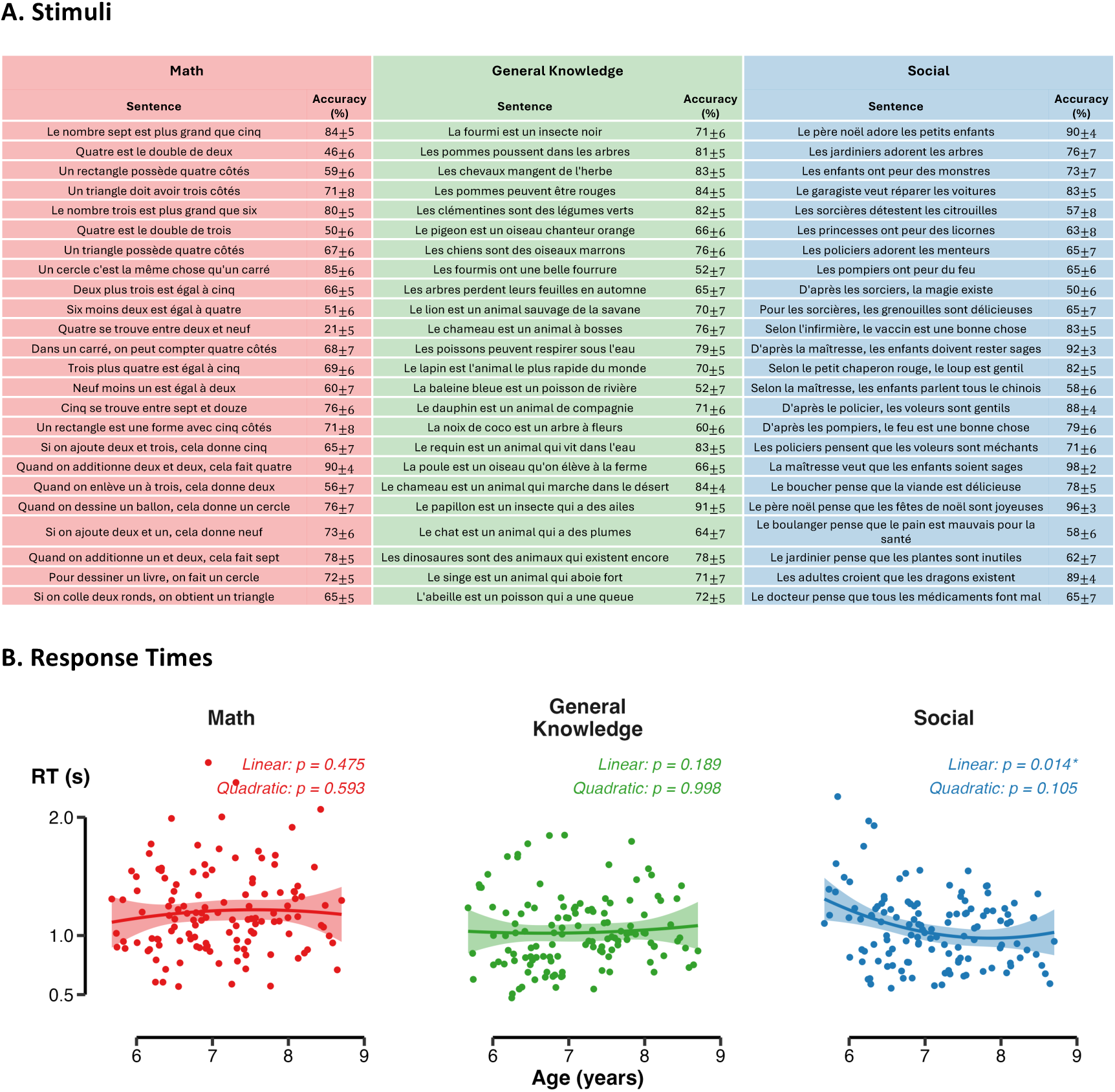
(A) Stimuli. French sentences used during the fMRI task, with 24 sentences covering different mathematical concepts, 24 of general knowledge and 24 of social reasoning. Different levels of complexity were included for each semantic condition. We reported the average accuracy and the standard error across participants and period for each individual sentence. **(B)** Reaction times (s) as a function of age’s participants, for correct trials only. A mixed-model regression was performed for each condition, with age as fixed effect and participant as random effect (*\*p < 0.05; **p < 0.01; ***p < 0.001*).

**Fig. S2.**
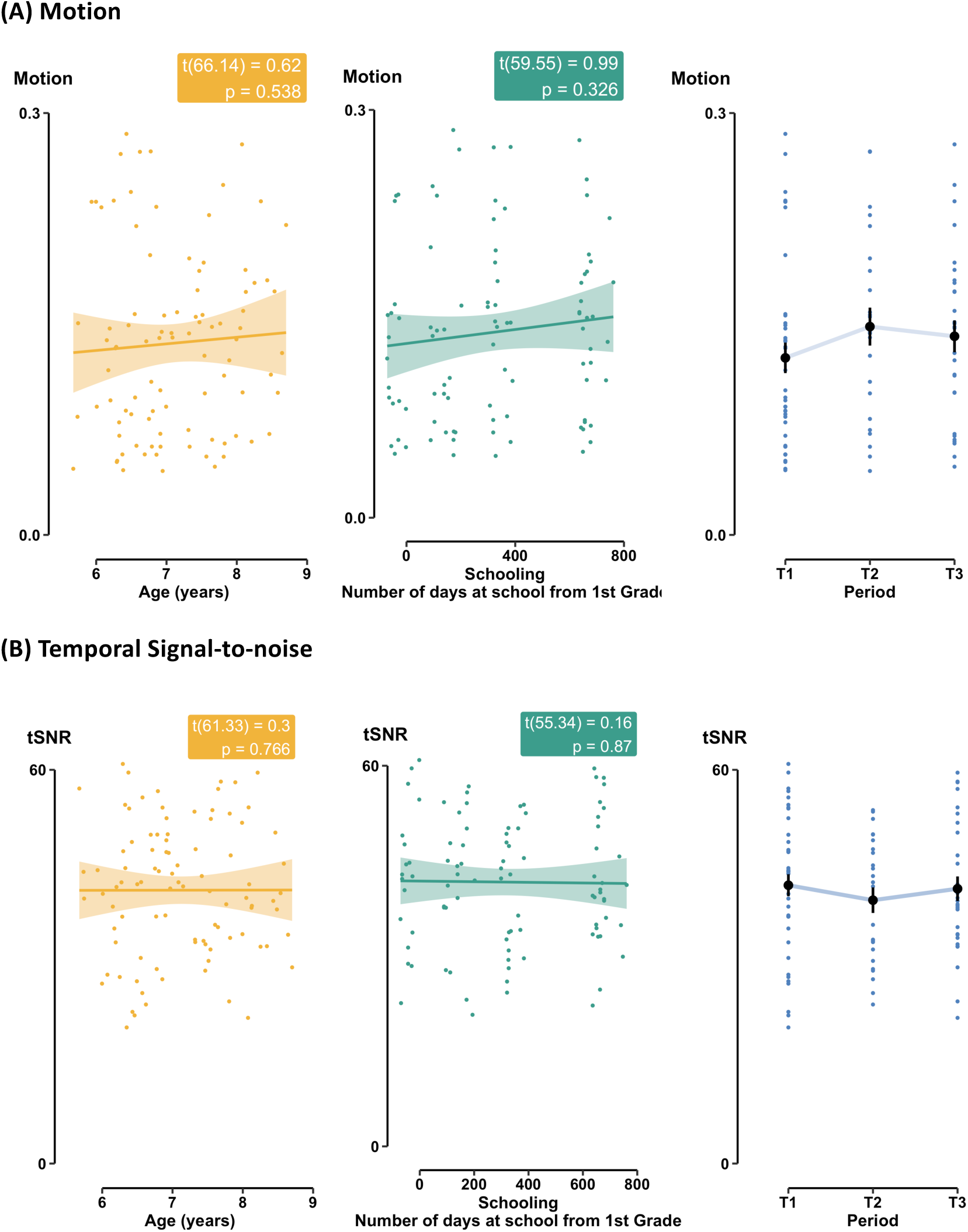
Data quality. **(A)** Overall amount of motion was computed for each included participant, averaged across axes and runs. Mixed-model regressions were performed with either age, number of days spent at school, or period as fixed effect, and participant as random effect (*\*p < 0.05; **p < 0.01; ***p < 0.001*). **(B)** Mean temporal signal-to-noise ratio (tSNR), averaged across voxels, were computed for each included participant. Mixed-model regressions were performed with either age, number of days spent at school, or period as fixed effect, and participant as random effect (*\*p < 0.05; **p < 0.01; ***p < 0.001*).

**Fig. S3.**
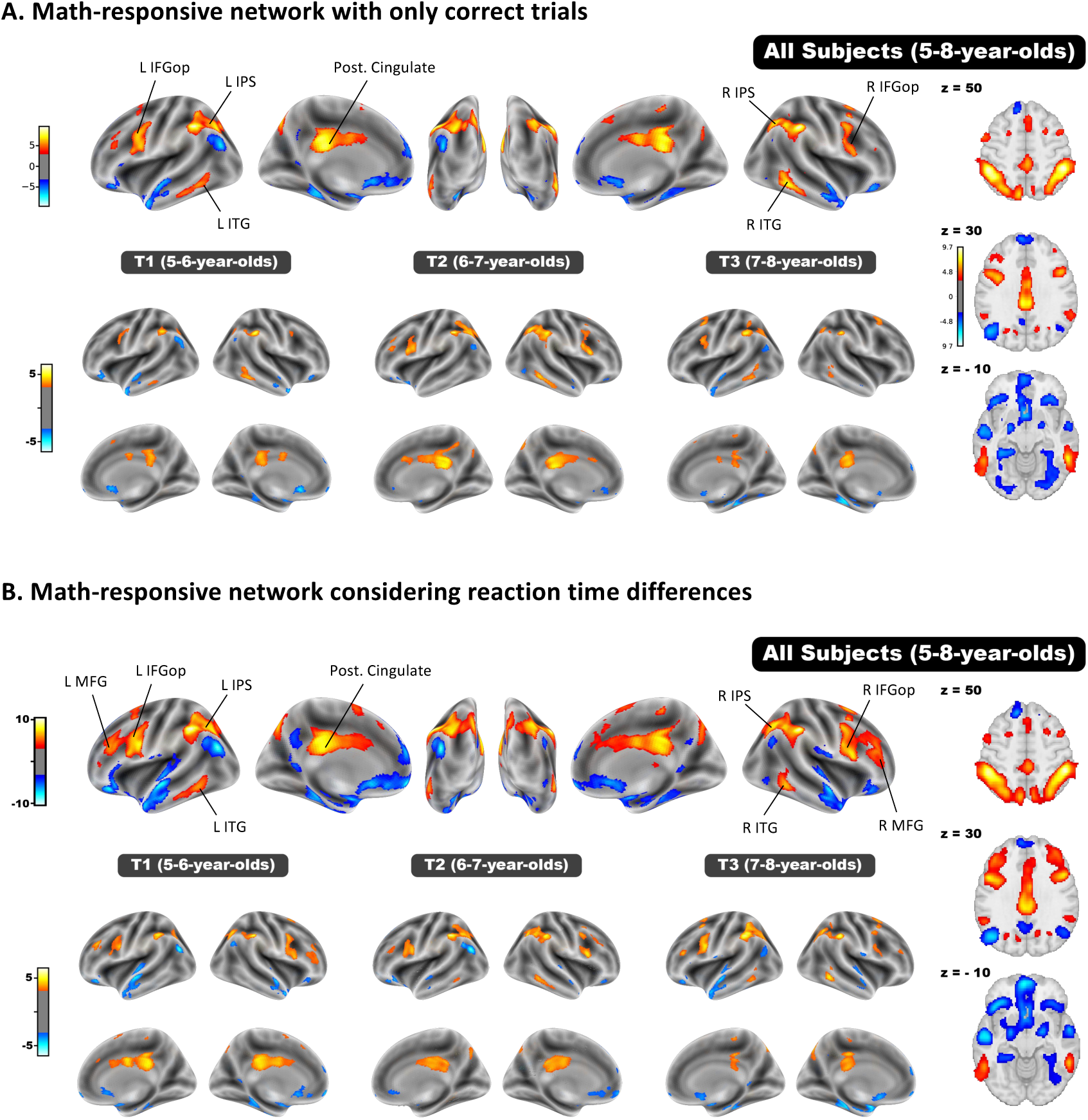
Control analysis for figure 2, in the same format. **(A)** Controlling for differences in accuracy across conditions: fMRI contrast between math and non-math sentences, restricted to trials correctly judged by children. **(B)** Controlling for differences in response time across conditions: fMRI contrast between math and non-math sentences, including response time for each sentence as a first-level covariate.

**Fig. S4.**
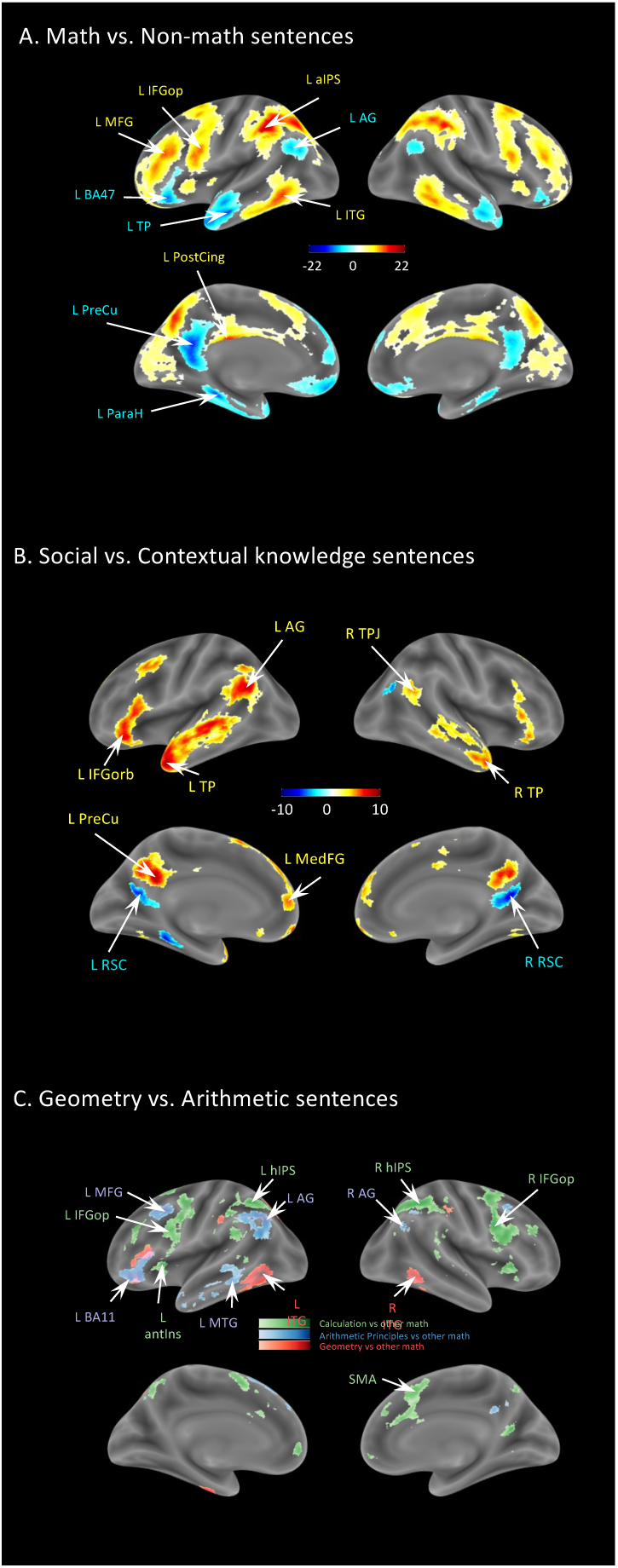
Adult networks from Moreno et al. (2025). Activation maps obtained in an independent cohort of adults, who were also asked to assess the veracity of sentences belonging to different semantic conditions. The sentences were not the same as those used in children. **(A)** Activation maps obtained for the contrast math vs. non-math sentences. **(B)** Activation maps obtained for the contrast social vs. contextual knowledge sentences (i.e., referring to a person vs. facts in a specific context). **(C)** Activation maps obtained for the contrast geometry vs. other mathematical sentences.

**Fig. S5.**
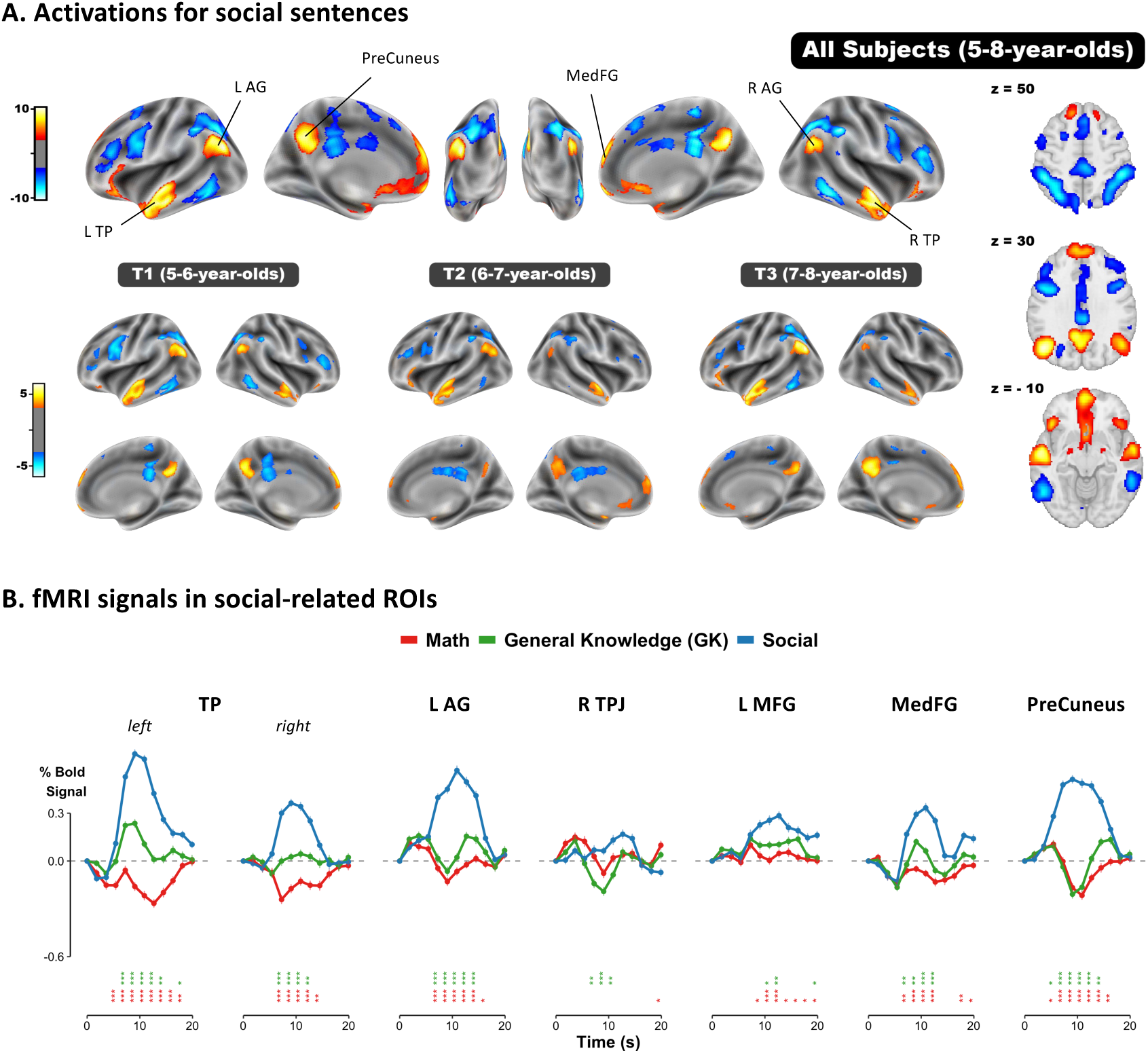
An adult-like social-responsive network in young children. **(A)** Activations obtained for the contrast between social and non-social sentences. The top row shows the joint analysis of all data, while bottom plots show separately the first fMRI (T1, 5-6-year-olds), second fMRI (T2, 6-7-year-olds), and third fMRI (T3, 7-8-year-olds). Voxel-wise p < 0.001, FDR-corrected α < 0.05. **(B)** Average fMRI signals in social-related ROIs defined in the study presented in S4(*54*). Stars indicate the significance of independent mixed-model regressions conducted at each time step (TR), with condition as fixed effect and participant as random effect (*•p < 0.1; *p < 0.05; **p < 0.01; ***p < 0.001*). Red stars: Math vs. Social. Green stars: Math vs. general knowledge.

**Fig. S6.**
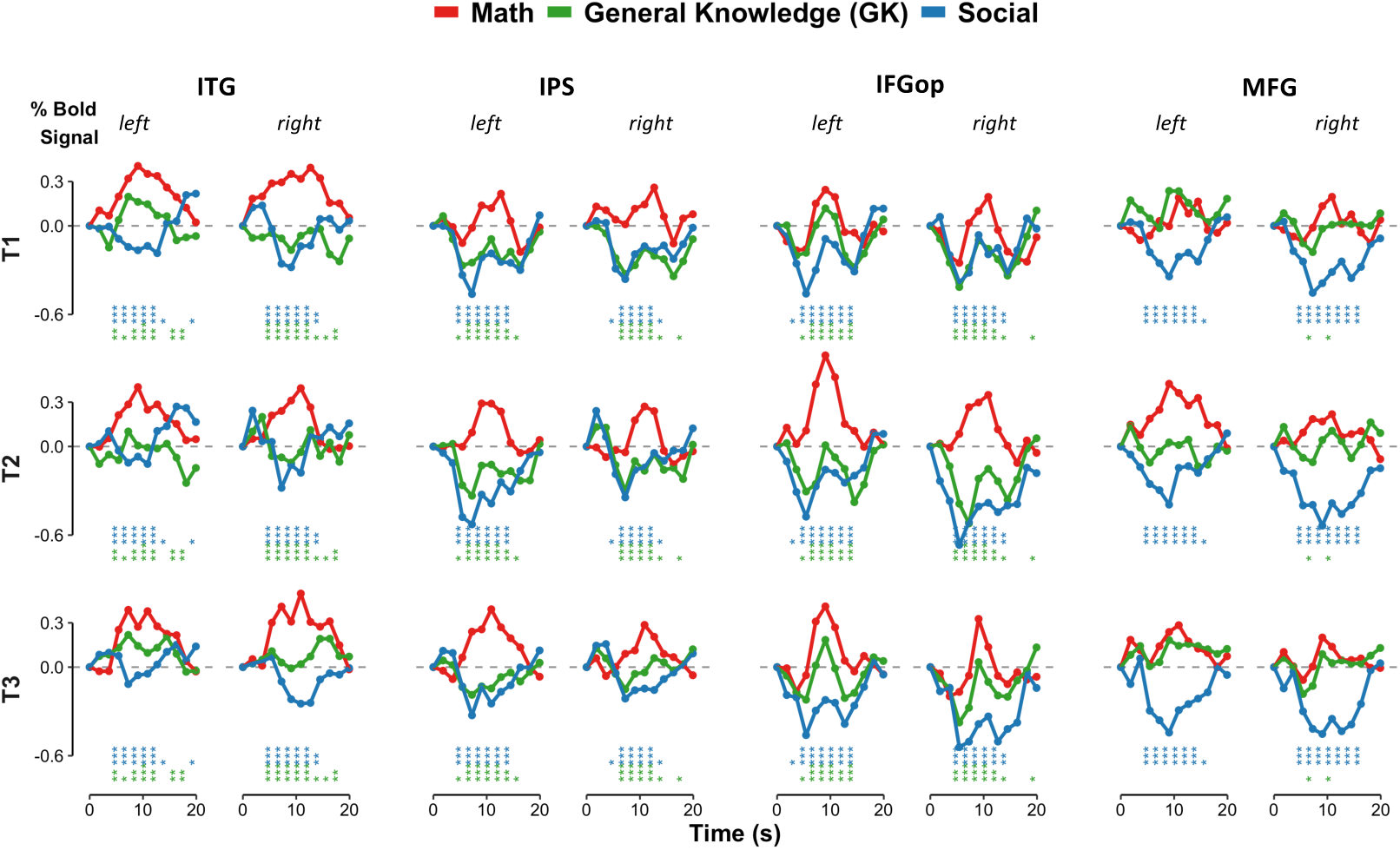
fMRI signals in adult math-responsive ROIs at each period. Average fMRI signals in math-responsive ROIs defined in the study presented in S4(*54*), at each period (T1, T2, and T3). Stars indicate the significance of independent mixed-model regressions conducted at each time step (TR), with condition as fixed effect and participant as random effect (*•p < 0.1; *p < 0.05; **p < 0.01; ***p < 0.001*). Red stars: Math vs. Social. Green stars: Math vs. general knowledge.

**Fig. S7.**
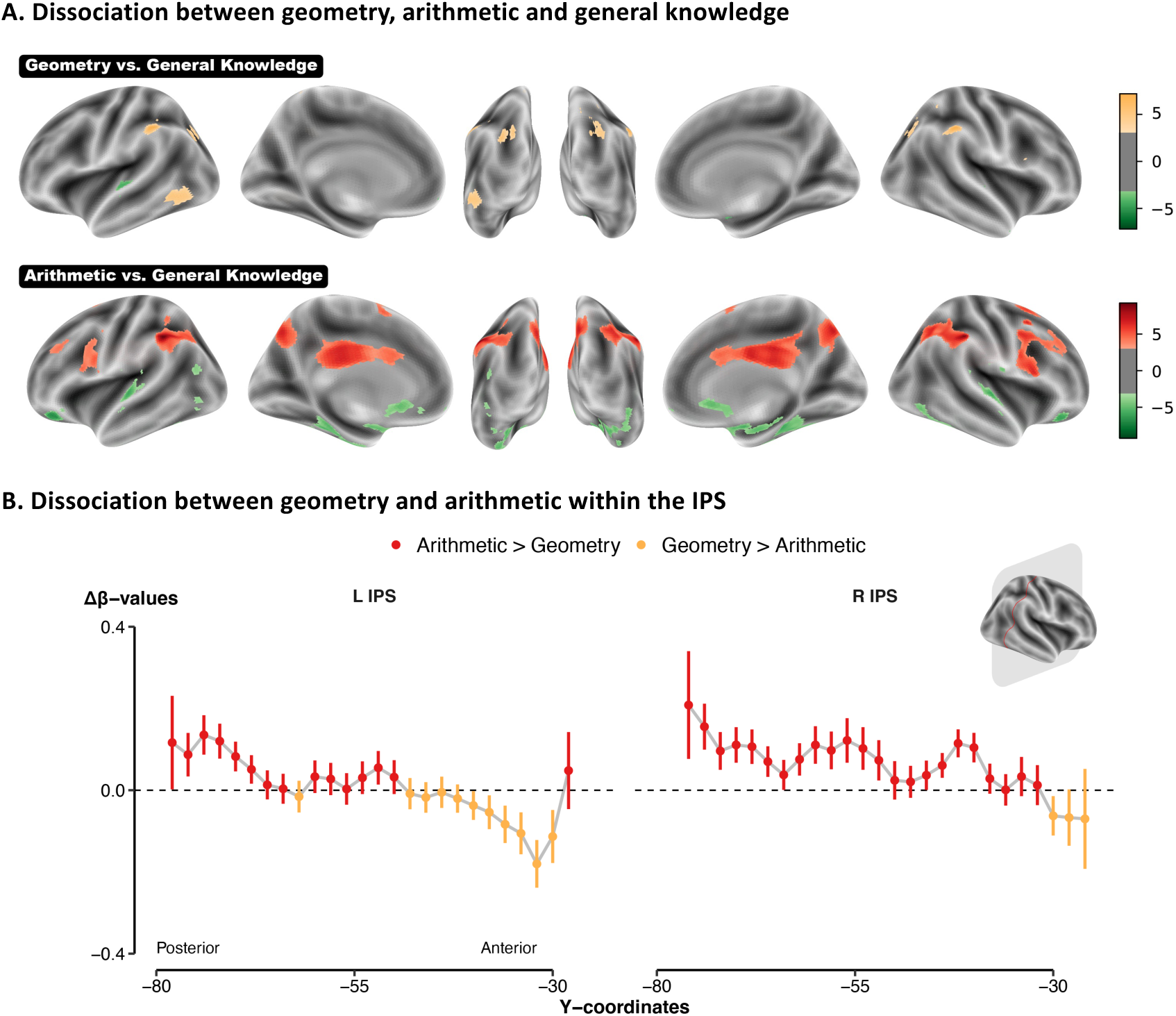
Geometric and arithmetic activations. **(A)** Activations obtained for the contrast geometry vs. general knowledge sentences, and arithmetic vs. general knowledge (voxel-wise p < 0.001, FDR-corrected α < 0.05). **(B)** Δ*β*-values (*β_Arithmetic_* – *β_Geometry_*) as a function of the y-coordinates in the left and right IPS. Mixed-model regressions with y-coordinates as fixed effect and participant as a random effect.

**Fig. S8.**
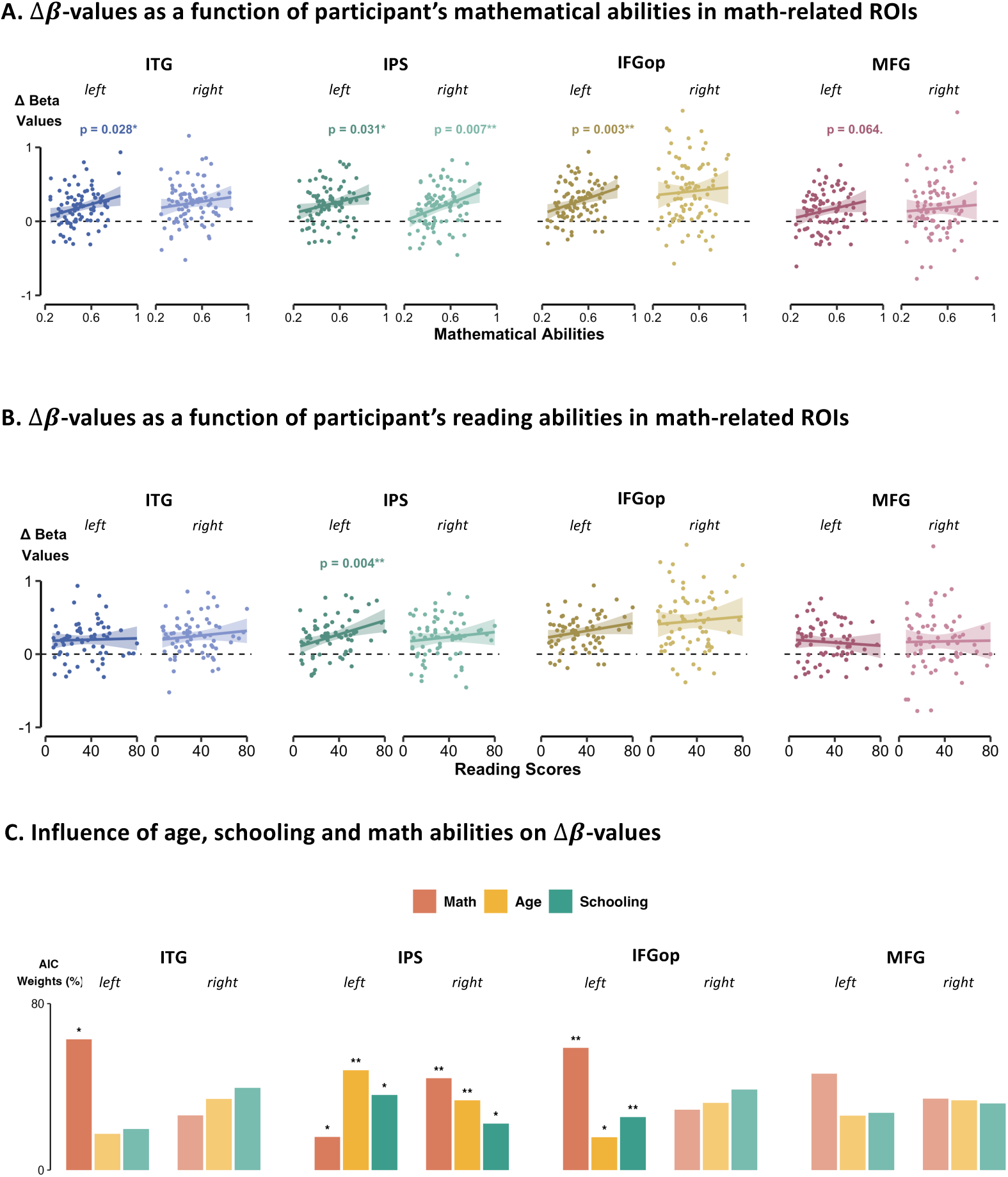
Partialing out the impact of age, schooling, and individual abilities. Each point represents the Δ*β*-values for each participant at each scan and for condition, within each ROI, as a function of children’s mathematical ability in **(A)** and as a function of children’s reading abilities in **(B).** Mixed-model regressions were conducted within each ROI, using individual abilities as fixed effect and participant as a random effect (*\*p < 0.05; **p < 0.01; ***p < 0.001*). **(C)** To disentangle the relative influences of these three developmental proxies, we compared a series of mixed models, each including a single predictor as fixed effect, and participant as random effect. Model fit was evaluated using the Akaike information criterion (AIC). To assess the robustness of model differences, we computed the Akaike weights, which estimates the probability that a given model is the best model, considering the evidence from all other models. Akaike weights (%) for each model computed within each ROI are represented here.

**Fig. S9.**
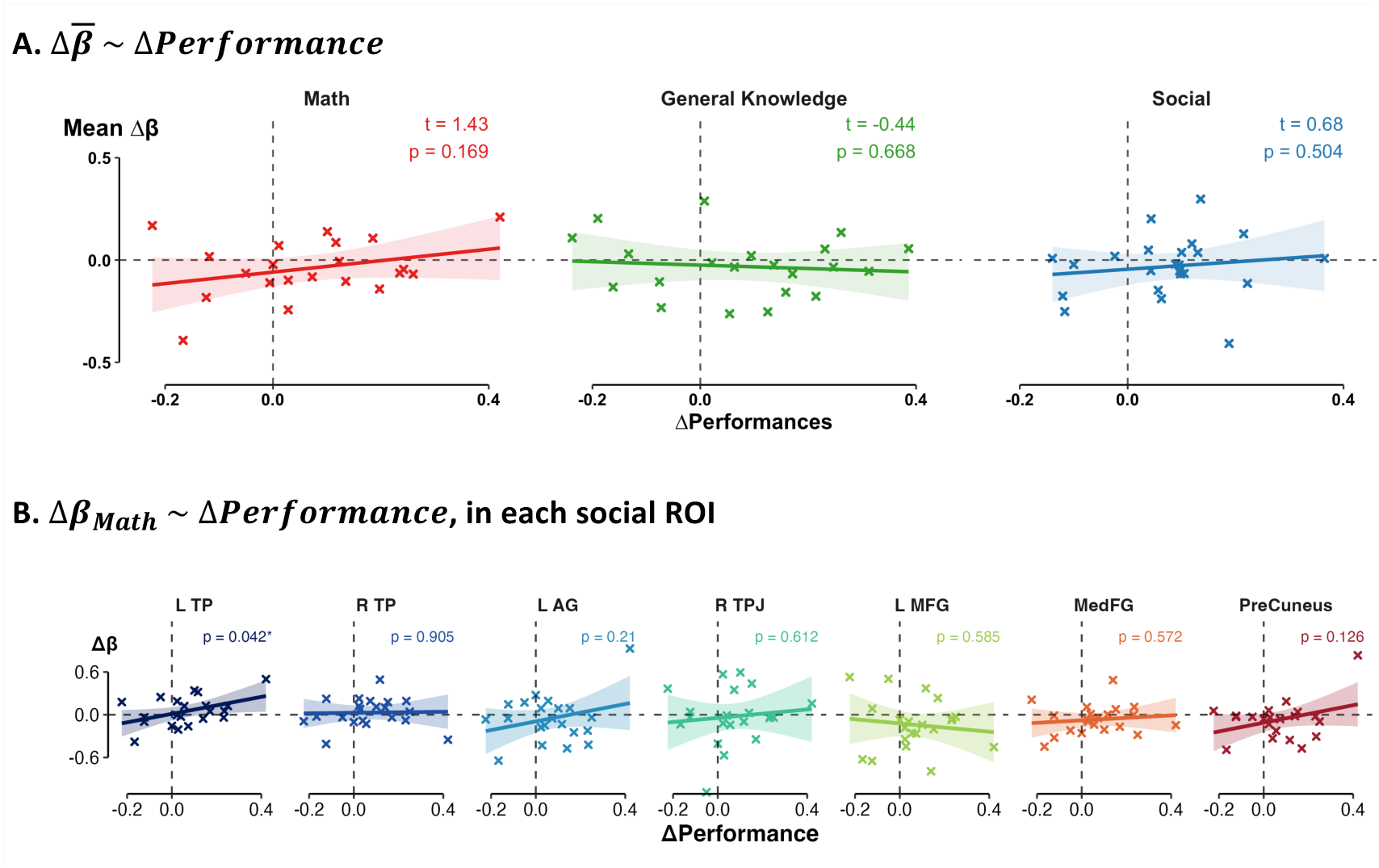
Control analyses for figure 6, in social-responsive ROIs. Only children scanned both at T1 and T3 (N = 26) were included in the analyses. For each child and sentence, we computed the change in brain activation (Δ*β*-values) by subtracting the *β*-values at T3 from those at T1, within each social-related ROI. For each sentence, we also computed the average behavioral performance across children at both T1 and T3 and used the difference (Δ*Perf*) as a proxy for change in comprehension. **(A)** Linear regressions were conducted for each condition, on mean Δ*β*-values averaged across children and social ROIs (*\*p < 0.05; **p < 0.01; ***p < 0.001*). **(B)** Linear regressions were performed for only math sentences, within each region-of-interest on mean Δ*β*-values averaged across children (*\*p < 0.05; **p < 0.01; ***p < 0.001*).

**Fig. S10.**
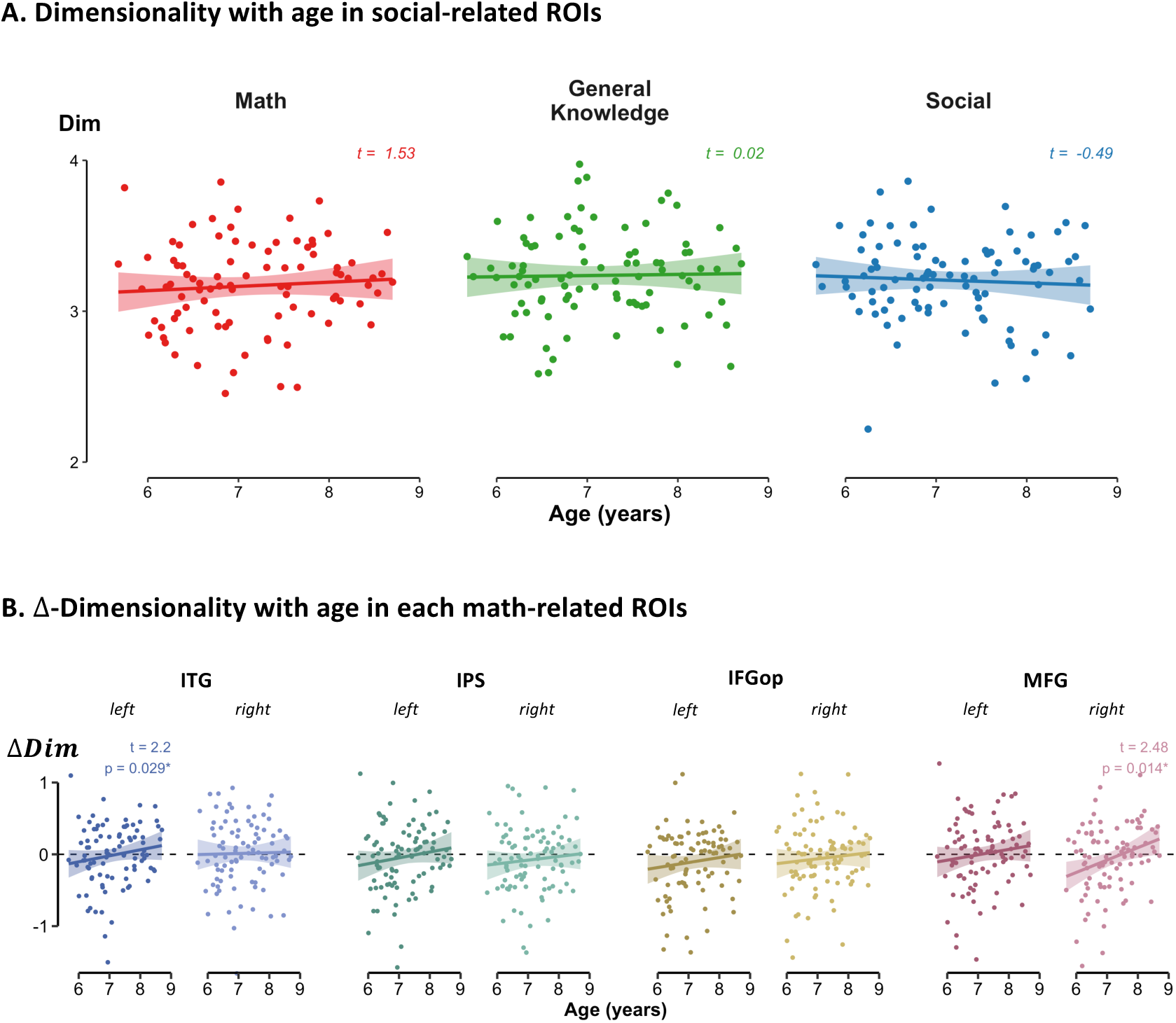
(A) No increase of neural space for mathematical concepts within social-responsive regions. Each point represents the dimensionality values for each participant and condition, at each scan, averaged across social-responsive ROIs. Mixed model regressions were conducted, using age as fixed effect, and including random intercepts and slopes for age per participant, and a random intercept for ROI (*\*p < 0.05; **p < 0.01; ***p < 0.001*). **(B)** Each point represents the Δ-dimensionality values (between math and non-math sentences) for each participant, at each scan and within each math-responsive ROI. Mixed model regressions were conducted within each ROI, using age as fixed effect, and participant as random effect (*\*p < 0.05; **p < 0.01; ***p < 0.001*).

## Supplementary tables

**Table S1.** Mixed-model coefficients for error rates across conditions. Coefficients obtained from a binomial mixed-model regression on error rates: *Error ∼ Condition × Age + (1|Subject)*, with condition being either math (reference condition), general knowledge or social. The predictor Age was standardized prior to the regression (GK = general knowledge).

|  | Estimate | Std. Error | Z-value | P-value |
| --- | --- | --- | --- | --- |
| (Intercept) | 0.68 | 0.091 | 7.34 |  |
| **GK vs. Math** | **-0.35** | **0.085** | **-4.08** | **< 0.001 (\*\*\*)** |
| **Social vs. Math** | **-0.45** | **0.086** | **-5.29** | **< 0.001 (\*\*\*)** |
| **Age** | **-0.24** | **0.061** | **-3.91** | **< 0.001 (\*\*\*)** |
| GK vs. Math x Age | 0.034 | 0.084 | 0.39 | 0.69 |
| Social vs. Math x Age | 0.029 | 0.085 | 0.33 | 0.74 |

**Table S2.** Mixed-model coefficients for reaction times across conditions. Coefficients obtained from a linear mixed-model regression on reaction times: *RT ∼ Condition × Age + (1|Subject)*, with condition being either math (reference condition), general knowledge or social. The predictor Age was standardized prior to the regression (GK = general knowledge).

|  | Estimate | Std. Error | DF | t-value | P-value |
| --- | --- | --- | --- | --- | --- |
| (Intercept) | 1.20 | 0.028 | 86.66 | 43.01 | < 0.001 (***) |
| **GK vs. Math** | **-0.14** | **0.022** | **3904.47** | **-6.63** | **< 0.001 (\*\*\*)** |
| **Social vs. Math** | **-0.14** | **0.022** | **3903.95** | **-6.29** | **< 0.001 (\*\*\*)** |
| Age | 0.014 | 0.017 | 3916.90 | 0.87 | 0.38 |
| GK vs. Math x Age | -0.013 | 0.022 | 3904.23 | -0.58 | 0.57 |
| **Social vs. Math x Age** | **-0.046** | **0.022** | **3903.65** | **-2.13** | **0.034 (\*)** |

**Table S3.** Geometry vs. arithmetic comparisons in math-responsive ROIs. Coefficients obtained from both frequentists mixed-model regressions and Bayesian ones, performed in each mathematical region-of-interest: β ∼ Condition + (1|Subject), with condition being either geometry (reference condition) or arithmetic. ROPE = Region of Practical equivalence (% of chance that BF_10_ is within [-0.1,0.1]). High %ROPE values indicate that the effect is likely negligible, whereas low values indicate that a meaningful difference between arithmetic and geometry is plausible. Although some regions showed uncorrected p-values below .05 (Left IFGop, Right MFG), none of these effects survived FDR correction (all pFDR ≥ .16). Bayesian analyses were consistent with this conclusion: most ROPE percentages exceeded 90%, indicating practically negligible effects. Only left and right IFGop and right MFG showed lower %ROPE values (< 50%), suggesting that small effects cannot be ruled out, although the Bayes Factors indicated only weak evidence.

| Region | Estimate | Std. Error | DF | t-value | p-value | p<sub>FDR</sub> | BF<sub>10</sub> | %ROPE |
| --- | --- | --- | --- | --- | --- | --- | --- | --- |
| Left ITG | -0.083 | 0.044 | 317.55 | -1.88 | 0.061 | 0.163 | 0.24 | 65.97 |
| Right ITG | -0.023 | 0.058 | 318.20 | -0.40 | 0.692 | 0.791 | 0.06 | 92.14 |
| Left IPS | -0.029 | 0.046 | 318.73 | -0.63 | 0.527 | 0.703 | 0.06 | 93.47 |
| Right IPS | 0.071 | 0.047 | 318.80 | 1.52 | 0.129 | 0.206 | 0.15 | 74.82 |
| Left IFGop | 0.103 | 0.049 | 315.68 | 2.12 | 0.035 (*) | 0.163 | 0.42 | 47.49 |
| Right IFGop | 0.132 | 0.081 | 319.37 | 1.64 | 0.101 | 0.202 | 0.27 | 35.19 |
| Left MFG | -0.009 | 0.050 | 315.89 | -0.18 | 0.858 | 0.858 | 0.05 | 97.70 |
| Right MFG | 0.112 | 0.056 | 318.1 | 2.01 | 0.045 (*) | 0.163 | 0.37 | 41.59 |

**Table S4.** Mixed-model coefficients for age effects in math-responsive ROIs. Coefficients obtained from mixed-model regressions performed in each mathematical region-of-interest: Δβ ∼ poly(Age, 2) + (1|Subject), with Δβ-values the difference between the activation elicited by mathematical sentences and the mean activation elicited by non-mathematical sentences (general knowledge and social).

| Region | Predictor | Estimate | Std. Error | DF | t-value | p-value | p<sub>FDR</sub> |
| --- | --- | --- | --- | --- | --- | --- | --- |
| Left ITG | Linear | 0.40 | 0.22 | 64.03 | 1.79 | 0.078 | 0.16 |
|  | Quadratic | -0.038 | 0.23 | 72.28 | -0.17 | 0.87 | 0.99 |
| Right ITG | Linear | 0.43 | 0.26 | 68.94 | 1.64 | 0.105 | 0.13 |
|  | Quadratic | -0.085 | 0.27 | 76.19 | -0.32 | 0.75 | 0.99 |
| **Left IPS** | **Linear** | **0.60** | **0.24** | **72.06** | **2.52** | **0.014 (\*)** | **0.037 (\*)** |
|  | Quadratic | -0.13 | 0.24 | 78.28 | -0.53 | 0.60 | 0.99 |
| **Right IPS** | **Linear** | **0.67** | **0.24** | **62.80** | **2.83** | **0.006 (\*\*)** | **0.032 (\*)** |
|  | Quadratic | -0.099 | 0.24 | 71.19 | -0.41 | 0.68 | 0.99 |
| **Left IFGop** | **Linear** | **0.56** | **0.21** | **59.45** | **2.73** | **0.008 (\*\*)** | **0.032 (\*)** |
|  | Quadratic | 0.037 | 0.21 | 67.57 | 0.17 | 0.86 | 0.99 |
| Right IFGop | Linear | 0.36 | 0.37 | 67.66 | 0.97 | 0.33 | 0.38 |
|  | Quadratic | -0.063 | 0.37 | 74.87 | -0.17 | 0.87 | 0.99 |
| Left MFG | Linear | 0.41 | 0.26 | 68.93 | 1.57 | 0.12 | 0.16 |
|  | Quadratic | 0.000 | 0.26 | 76.20 | -0.002 | 0.10 | 0.99 |
| Right MFG | Linear | 0.16 | 0.34 | 63.60 | 0.45 | 0.65 | 0.65 |
|  | Quadratic | 0.051 | 0.35 | 72.19 | 0.15 | 0.88 | 0.99 |

**Supplementary table S5.**
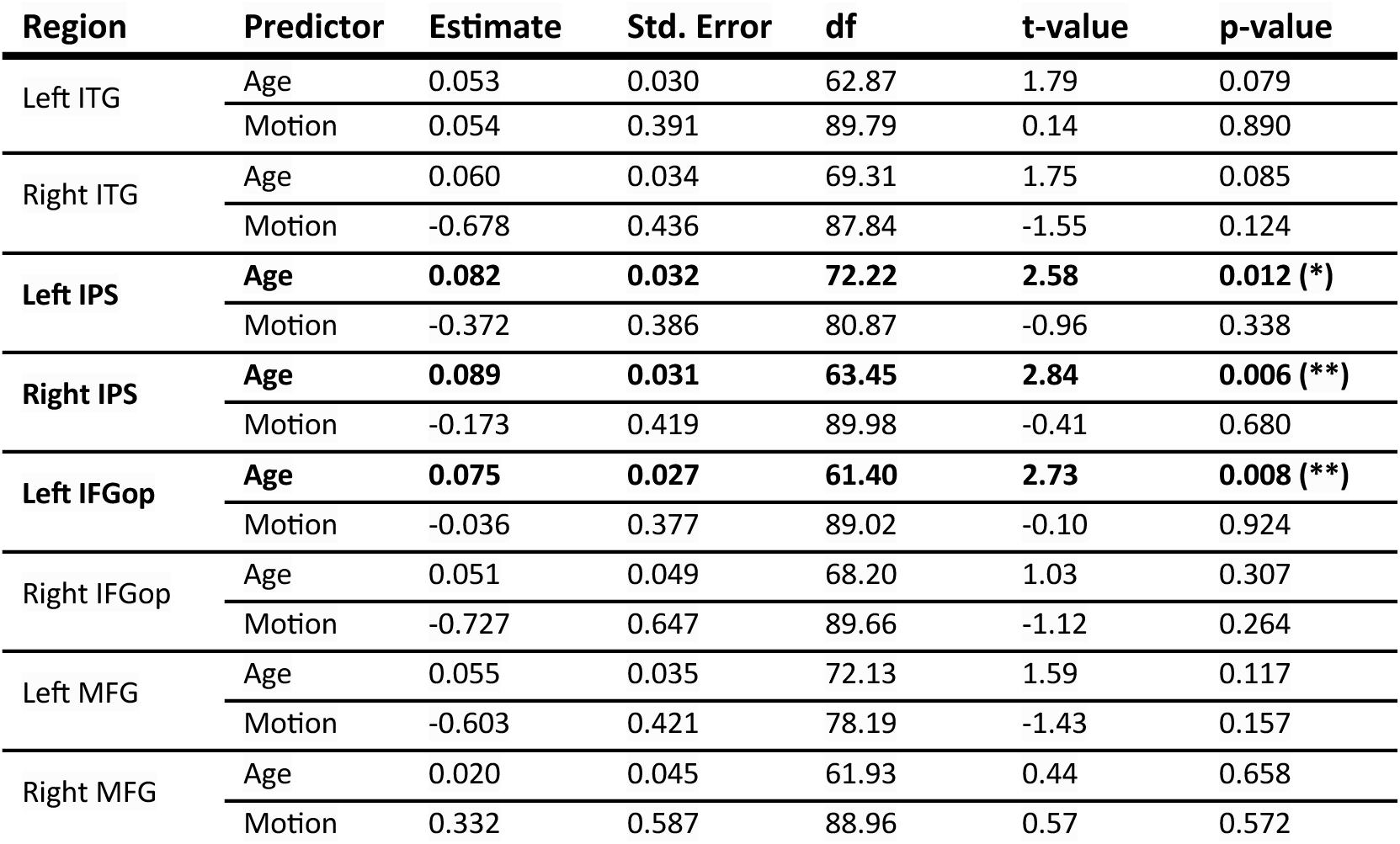
Controlling for motion effects in developmental changes in math ROIs. Coefficients obtained from mixed-model regressions performed in each mathematical region-of-interest: Δβ ∼ Age + Motion + (1|Subject), with Δβ-values the difference between the activation elicited by mathematical sentences and the mean activation elicited by non-mathematical sentences (general knowledge and social).

**Table S6.** Controlling for noise effects in developmental changes in math ROIs. Coefficients obtained from mixed-model regressions performed in each mathematical region-of-interest: Δβ ∼ Age + tSNR + (1|Subject), with Δβ-values the difference between the activation elicited by mathematical sentences and the mean activation elicited by non-mathematical sentences (general knowledge and social).

| Region | Predictor | Estimate | Std. error | df | t-value | p-value |
| --- | --- | --- | --- | --- | --- | --- |
| Left ITG | Age | 0.052 | 0.030 | 62.51 | 1.77 | 0.082 |
|  | tSNR | 0.002 | 0.003 | 86.88 | 0.76 | 0.452 |
| Right ITG | Age | 0.055 | 0.033 | 70.27 | 1.65 | 0.103 |
|  | tSNR | **0.008** | **0.003** | **83.49** | **2.85** | **0.006 (\*\*)** |
| Left IPS | Age | **0.080** | **0.031** | **72.24** | **2.53** | **0.014 (\*)** |
|  | tSNR | 0.004 | 0.002 | 73.98 | 1.73 | 0.087 |
| Right IPS | Age | **0.088** | **0.031** | **63.67** | **2.82** | **0.006 (\*\*)** |
|  | tSNR | 0.003 | 0.003 | 88.66 | 1.19 | 0.237 |
| Left IFGop | Age | **0.072** | **0.027** | **63.47** | **2.61** | **0.011 (\*)** |
|  | tSNR | 0.005 | 0.002 | 89.51 | 1.94 | 0.056 |
| Right IFGop | Age | 0.045 | 0.049 | 69.20 | 0.93 | 0.358 |
|  | tSNR | 0.008 | 0.004 | 87.55 | 1.98 | 0.051 |
| Left MFG | Age | 0.052 | 0.034 | 90.00 | 1.51 | 0.135 |
|  | tSNR | 0.006 | 0.003 | 90.00 | 2.19 | 0.031 |
| Right MFG | Age | 0.022 | 0.045 | 62.62 | 0.49 | 0.629 |
|  | tSNR | - | 0.004 | 86.30 | - | 0.608 |

**Table S7.**
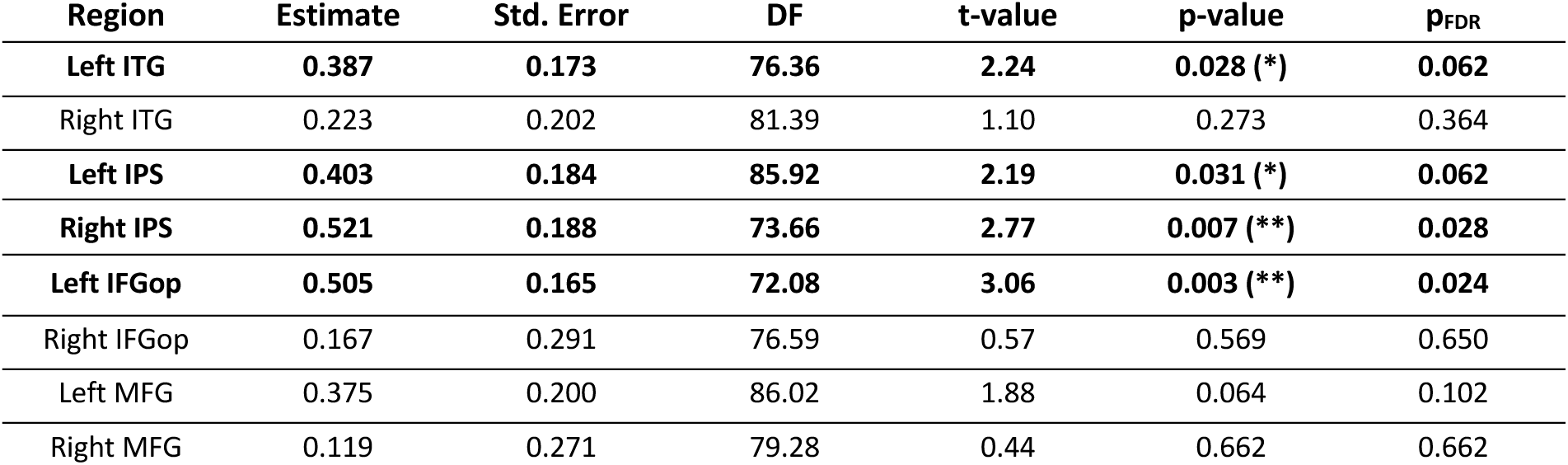
Math-ability effects in math-responsive ROIs. Coefficients obtained from mixed-model regressions performed in each mathematical region-of-interest: Δβ ∼ Math, with Δβ-values the difference between the activation elicited by mathematical sentences and the mean activation elicited by non-mathematical sentences (general knowledge and social), and math the individual mathematical abilities obtained from the number screener test.

| Region | Estimate | Std. Error | DF | t-value | p-value | p<sub>FDR</sub> |
| --- | --- | --- | --- | --- | --- | --- |
| **Left ITG** | **0.387** | **0.173** | **76.36** | **2.24** | **0.028 (\*)** | **0.062** |
| Right ITG | 0.223 | 0.202 | 81.39 | 1.10 | 0.273 | 0.364 |
| **Left IPS** | **0.403** | **0.184** | **85.92** | **2.19** | **0.031 (\*)** | **0.062** |
| **Right IPS** | **0.521** | **0.188** | **73.66** | **2.77** | **0.007 (\*\*)** | **0.028** |
| **Left IFGop** | **0.505** | **0.165** | **72.08** | **3.06** | **0.003 (\*\*)** | **0.024** |
| Right IFGop | 0.167 | 0.291 | 76.59 | 0.57 | 0.569 | 0.650 |
| Left MFG | 0.375 | 0.200 | 86.02 | 1.88 | 0.064 | 0.102 |
| Right MFG | 0.119 | 0.271 | 79.28 | 0.44 | 0.662 | 0.662 |

**Table S8.** Reading-fluency effects in math-responsive ROIs. Coefficients obtained from mixed-model regressions performed in each mathematical region-of-interest: Δβ ∼ Reading, with Δβ-values the difference between the activation elicited by mathematical sentences and the mean activation elicited by non-mathematical sentences (general knowledge and social), and reading the individual reading fluency (number of words read in one min).

| Region | Estimate | Std. Error | DF | t-value | p-value | p<sub>FDR</sub> |
| --- | --- | --- | --- | --- | --- | --- |
| Left ITG | 0.001078 | 0.001611 | 66.270 | 0.669 | 0.506 | 0.720 |
| Right ITG | 0.002576 | 0.001702 | 66.904 | 1.513 | 0.135 | 0.448 |
| **Left IPS** | **0.004734** | **0.001593** | **67.000** | **2.972** | **0.004 (\*\*)** | **0.032** |
| Right IPS | 0.002007 | 0.001848 | 66.809 | 1.086 | 0.281 | 0.562 |
| Left IFGop | 0.002146 | 0.001541 | 66.873 | 1.392 | 0.168 | 0.448 |
| Right IFGop | 0.001107 | 0.002687 | 66.749 | 0.412 | 0.682 | 0.779 |
| Left MFG | -0.001069 | 0.001734 | 67.000 | -0.616 | 0.540 | 0.720 |
| Right MFG | 0.000268 | 0.002584 | 67.000 | 0.104 | 0.918 | 0.918 |

**Table S9.** Amplified vs. reduced voxels percentages in math-responsive ROIs. Coefficients obtained from linear regressions performed in each mathematical region-of-interest: Percentages ∼ Category, comparing the percentages of amplified voxels (Δ*β*(*T*_3_) > Δ*β*(*T*_1_), and Δ*β*(*T*_1_) > 0) vs. reduced voxels (Δ*β*(*T*_3_) < Δ*β*(*T*_1_), and Δ*β*(*T*_1_) > 0). Degree of freedom = 25.

| Region | Estimate | Std. Error | t | p-value | p<sub>FDR</sub> |
| --- | --- | --- | --- | --- | --- |
| Left ITG | 1.70 | 2.32 | 0.73 | 0.469 | 0.536 |
| Right ITG | 6.52 | 2.94 | 2.22 | 0.0313 (*) | 0.0682 |
| Left IPS | 5.95 | 2.04 | 2.92 | 0.0052 (**) | 0.0418 (*) |
| Right IPS | 4.98 | 2.28 | 2.18 | 0.0341 (*) | 0.0682 |
| Left IFGop | 6.13 | 2.34 | 2.62 | 0.0115 (*) | 0.0461 (*) |
| Right IFGop | 4.73 | 4.60 | 1.03 | 0.309 | 0.412 |
| Left MFG | 4.65 | 2.38 | 1.96 | 0.0558 | 0.0893 |
| Right MFG | 0.36 | 3.30 | 0.11 | 0.913 | 0.913 |

**Table S10.** Recruited vs. dropout voxel percentages in math-responsive ROIs. Coefficients obtained from linear regressions performed in each mathematical region-of-interest: Percentages ∼ Category, comparing the percentage of recruited voxels (Δ*β*(*T*_3_) > 0, and Δ*β*(*T*_1_) < 0) vs. dropout voxels (Δ*β*(*T*_3_) < 0, and Δ*β*(*T*_1_) > 0). Degree of freedom = 25.

| Region | Estimate | Std. Error | t | p-Value | p<sub>FDR</sub> |
| --- | --- | --- | --- | --- | --- |
| Left ITG | 5.06 | 2.18 | 2.32 | 0.0245 (*) | 0.0655 |
| Right ITG | 1.00 | 2.87 | 0.35 | 0.728 | 0.728 |
| Left IPS | 6.40 | 2.52 | 2.54 | 0.0141 (*) | 0.0564 |
| Right IPS | 4.93 | 2.27 | 2.17 | 0.0348 (*) | 0.0696 |
| Left IFGop | 7.43 | 2.16 | 3.45 | 0.0012 (**) | 0.0093 (**) |
| Right IFGop | 2.14 | 4.01 | 0.53 | 0.595 | 0.681 |
| Left MFG | 5.42 | 2.92 | 1.85 | 0.0696 | 0.111 |
| Right MFG | 3.31 | 3.59 | 0.92 | 0.362 | 0.482 |

**Table S11.** Brain–performance change relationships in math-responsive ROIs for math sentences. Coefficients obtained from linear regressions performed in each mathematical region-of-interest: Δβ ∼ ΔPerf, with each data point corresponding to an individual mathematical sentence. Δβ-values correspond to the difference in brain activation between T3 and T1, elicited by each sentence and averaged across participants. ΔPerf represents the change in behavioral accuracy between T3 and T1 for each sentence, averaged across participants.

| Region | Estimate | Std. Error | DF | t-value | p-value | p<sub>FDR</sub> |
| --- | --- | --- | --- | --- | --- | --- |
| Left ITG | -0.93 | 0.39 | 19 | -2.35 | 0.030 (*) | 0.06 |
| Right ITG | -1.36 | 0.34 | 19 | -4.04 | 0.001 (***) | 0.008 (***) |
| Left IPS | -0.36 | 0.57 | 19 | -0.63 | 0.53 | 0.61 |
| Right IPS | -0.20 | 0.48 | 19 | -0.41 | 0.69 | 0.69 |
| Left IFGop | -1.01 | 0.42 | 19 | -2.38 | 0.028 (*) | 0.06 |
| Right IFGop | -1.13 | 0.58 | 19 | -1.95 | 0.066 | 0.11 |
| Left MFG | -0.69 | 0.49 | 19 | -1.41 | 0.17 | 0.23 |
| Right MFG | -1.06 | 0.40 | 19 | -2.73 | 0.013 (*) | 0.05 |

**Table S12.**
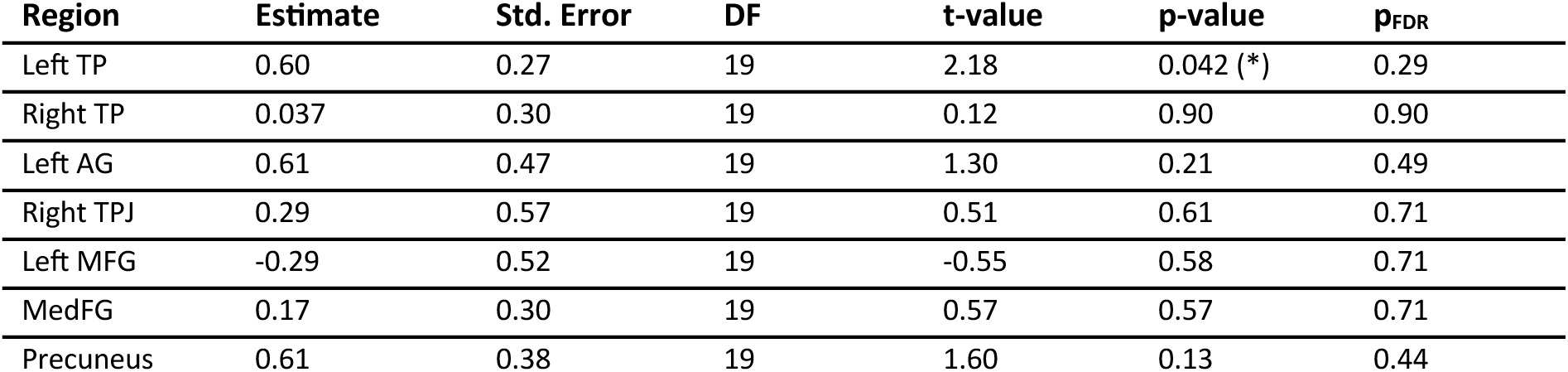
Brain–performance change relationships in social-responsive ROIs for math sentences. Coefficients obtained from linear regressions performed in each social region-of-interest: Δβ ∼ ΔPerf, with each data point corresponding to an individual mathematical sentence. Δβ-values correspond to the difference in brain activation at T3 minus T1, elicited by each sentence and averaged across participants. ΔPerf is the difference between average performance at T3 minus T1 for each sentence, averaged across participants.

**Table S13.**
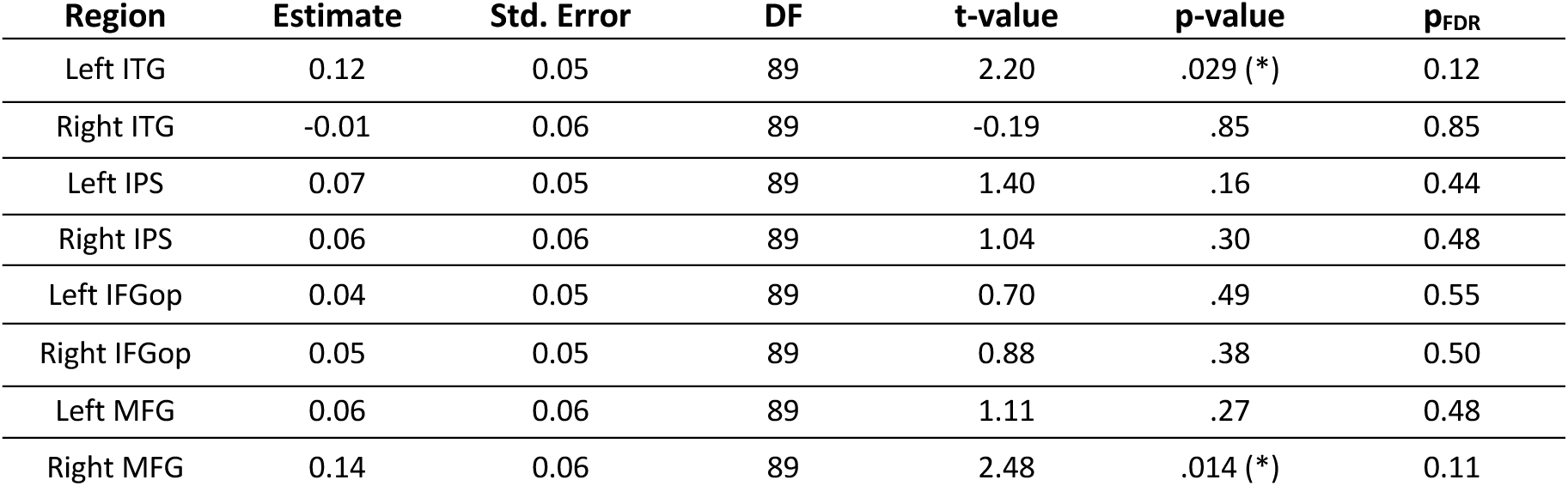
Dimensionality changes in math-responsive ROIs. Coefficients obtained from linear regressions performed in each math-responsive ROI: ΔDim ∼ Age. ΔDim-values correspond to the difference between the dimensionality for mathematical sentences and the mean dimensionality for non-mathematical sentences (general knowledge and social).

| Region | Estimate | Std. Error | DF | t-value | p-value | p<sub>FDR</sub> |
| --- | --- | --- | --- | --- | --- | --- |
| Left ITG | 0.12 | 0.05 | 89 | 2.20 | .029 (*) | 0.12 |
| Right ITG | -0.01 | 0.06 | 89 | -0.19 | .85 | 0.85 |
| Left IPS | 0.07 | 0.05 | 89 | 1.40 | .16 | 0.44 |
| Right IPS | 0.06 | 0.06 | 89 | 1.04 | .30 | 0.48 |
| Left IFGop | 0.04 | 0.05 | 89 | 0.70 | .49 | 0.55 |
| Right IFGop | 0.05 | 0.05 | 89 | 0.88 | .38 | 0.50 |
| Left MFG | 0.06 | 0.06 | 89 | 1.11 | .27 | 0.48 |
| Right MFG | 0.14 | 0.06 | 89 | 2.48 | .014 (*) | 0.11 |

